# Reversible ubiquitination of integrated domain controls paired NLR immune receptor complex homeostasis

**DOI:** 10.1101/2024.07.01.599856

**Authors:** Zhiyi Chen, Jianhua Huang, Jianyu Li, Frank L.H. Menke, Jonathan D.G. Jones, Hailong Guo

## Abstract

Plant intracellular NLR immune receptors can function individually or in pairs to detect pathogen effectors and activate immune responses. NLR homeostasis has to be tightly regulated to ensure proper defense without triggering autoimmunity. However, in contrast to singleton NLRs, the mechanisms controlling the paired NLRs complex homeostasis are less understood. The paired Arabidopsis RRS1/RPS4 immune receptor complex confers disease resistance through effector recognition mediated by the integrated WRKY domain of RRS1. Here, through proximity labelling, we reveal a ubiquitination-deubiquitination cycle that controls the homeostasis of the RRS1/RPS4 complex. E3 ligase RARE directly binds and ubiquitinates RRS1’s WRKY domain to promote its proteasomal degradation, thereby destabilizing RPS4 indirectly and compromising the stability and function of the RRS1/RPS4 complex. Conversely, the deubiquitinating enzymes UBP12/UBP13 deubiquitinate RRS1’s WRKY domain, counteracting RARE’s effects. Interestingly, the abundance of WRKY transcription factors WRKY70 and WRKY41 is also regulated by RARE and UBP12/UBP13. Phylogenetic analysis suggests this regulation likely transferred from WRKY70/WRKY41 to RRS1 upon WRKY domain integration. Our findings improve our understanding of homeostatic regulation of paired NLR complex and uncover a new paradigm whereby domain integration can co-opt preexisting post-translational modification to regulate novel protein functions.

## Introduction

Phytopathogens secrete effector proteins into plant cells to suppress plant immunity for colonization. To circumvent pathogen invasion, plants have evolved intracellular nucleotide-binding leucine-rich repeat immune receptors (NLRs) that recognize effectors, either by directly binding effectors or by monitoring effector-mediated modification of “guardees” or “decoys” to activate effector-triggered immunity (ETI). Activation of ETI involves calcium influx, global transcriptional reprogramming and programmed cell death called hypersensitive response (HR)^1,2^. Canonical NLRs contain a C-terminal leucine-rich repeat (LRR) domain, a central nucleotide-binding (NB) domain and a variable N-terminal domain including Toll/Interleukin-1 receptor (TIR) domain, coiled-coil (CC) domain, or Resistance to Powdery Mildew 8 (RPW8)-like CC (CC_R_) domain^3^. Based on their N-termini, NLRs are classified into TNLs (TIR-NLRs), CNLs (CC-NLRs) and RNLs (CC_R_-NLR). Upon effector recognition, both CNLs and TNLs oligomerize into complexes termed resistosomes that activate their calcium-permeable ion channel activity or NADase combined with ribosyl-transferase activities, respectively, with the latter producing small molecules that contribute to RNL activation^4–7^.

A subclass of NLRs carries additional noncanonical domain(s) that appear to have evolved by integrating authentic effector targets into the canonical NLR structure. These NLRs with extra domains are thus commonly referred to as NLRs with integrated domains (NLR-IDs) that account for around 10-15% of the NLRome of many plant species^8,9^. Several functionally characterized NLR-IDs require another genetically linked canonical NLR to work in pairs for their function and are therefore known as NLR pairs (or sensor-executor pairs)^3,10^. Sensor and executor NLRs form pre-activation complexes in the resting state and, upon binding of effectors to the integrated domains, sensor NLRs undergo a series of intra- and intermolecular conformational rearrangements that relay effector perception to the executor NLRs for immune signaling initiation^11–13^. The Arabidopsis RRS1/RPS4 immune receptor complex, one of the best-studied paired NLRs, confers resistance to bacterial pathogens *Pseudomonas syringae* pv. *pisi*. and *Ralstonia solanacearum* by recognizing effectors AvrRps4 and PopP2, respectively^14^. RRS1 from Arabidopsis accession Col-0 (termed RRS1-S) recognizes only AvrRps4, while the RRS1 from accessions Nd-1 and Ws-2 (termed RRS1-R), with a longer C-terminal extension beyond the end of the WRKY domain compared to RRS1-S, confers responsiveness to both AvrRps4 and PopP2^15,16^. AvrRps4 and PopP2 are both detected by the C-terminal integrated WRKY domain of RRS1-R that mimics the authentic targets of effectors and derepress the complex in distinct ways^17,18^. Whether the WRKY ID possesses additional roles in controlling immune activation and regulation of RRS1/RPS4 complex remains uncharacterized.

In recent years, it became apparent that NLRs are associated with NLR-interacting proteins to achieve a balanced level of immunity^19^. Ubiquitination, a reversible and dynamic process of tagging ubiquitin to a substrate protein, plays a pivotal role in maintaining NLRs homeostasis. A few E3 ligases were found to associate with NLRs to regulate their protein degradation via 26S proteasome. For example, the F-box containing E3 ligase CPR1 was shown to target NLR proteins SNC1 and RPS2 for degradation^20^. Two functionally redundant RING-type E3 ligases, MUSE1/MUSE2, regulate the turnover of SNC1’s partners, SIKIC1/2/3^21^. Interestingly, another two closely related RING-type E3 ligases, SNIPER1/SNIPER2, broadly regulate the homeostasis of diverse sensor NLRs including SNC1, SUMM2, RPP4 and RPM1^22^. However, all E3 ligases that have been identified to date are from the investigations of single NLR proteins. Furthermore, to date, no deubiquitinating enzymes have been identified that directly remove ubiquitin from NLRs and limit their 26S proteasomal degradation.

NLR-ID fusion proteins are likely formed through DNA transposition and/or ectopic recombination, the major contributors to domain shuffling^23^. Domain shuffling constantly occurred during the dynamic evolutionary history of NLRs and facilitated the diversity of NLR repertoires, allowing NLRs to cope with the rapid evolution of pathogen effectors. Fusion of effector-targeted domains into NLRs via domain shuffling creates NLR-IDs that can reveal the action of effectors^24,25^. As more paired NLR/NLR-ID complex have now been identified, how such NLR/NLR-ID complexes are regulated and how domain shuffling affects fusion proteins regulation remains unknown. In this study, using TurboID-based proximity labeling, we identified the E3 ligase RARE that associates with RRS1/RPS4 complex through directly interacts with RRS1 but not with RPS4. RARE targets RRS1 for ubiquitination through the WRKY domain and destabilizes the complex abundance in a RRS1-dependent manner, thus suppressing immune responses conditioned by RRS1/RPS4. Additionally, we found that two closely related deubiquitinating enzymes UBP12/UBP13 antagonize the action of RARE through deubiquitinating RRS1. Interestingly, RARE and UBP12/UBP13 also modulate the homeostasis of WRKY transcription factors WRKY70 and WRKY41. Phylogenetic analysis suggests that such regulation was most likely transferred from the WRKY transcription factors to RRS1 during WRKY domain integration. Thus, our findings not only uncover the reversible ubiquitination of ID in regulating the homeostasis of paired NLR/NLR-ID complex, but also reveal a new paradigm whereby domain integration can transfer pre-existing post-translational regulatory mechanisms like ubiquitination to regulate novel protein functions.

## Results

### Identification of RARE and UBP12/13 in the proxiomes of RRS1

To unravel the regulation and signaling of RRS1/RPS4 immune receptor complex, we employed proximity labeling to identify their interactors^26^. Both RRS1 and RPS4 were fused with highly active biotin ligase TurboID tagged with a FLAG epitope for proximity labeling. In biotinylation assays, RRS1-TurboID induced more pronounced biotinylation compared to RRS4-TurboID in *Nicotiana benthamiana* (*N. benthamiana*), likely due to its higher protein abundance (Figure S1A). Alongside cognate effectors AvrRps4 and PopP2, co-expression of RRS1 and RPS4 triggers HR in *N. tabacum*. However, the fusion of the TurboID tag impaired the HR only when fused to RPS4 but not RRS1(Figure 1A), suggesting that RRS1-TurboID, but not RPS4-TurboID, is fully functional. Consistently, resistance against *Pseudomonas syringae* (*Pst*) DC3000 carrying AvrRps4 was restored in Col-0 *rrs1arrs1b* mutants complemented with the RRS1-TurboID transgene driven by pAt2 promoter (Figure 1B and C), which allows moderate constitutive expression^27,28^. We therefore utilized RRS1-R-TurboID for further proximity labeling assays since RRS1-S is likely a derived allele from RRS1-R as a result of premature stop codon^15,16^.

**Figure 1.**
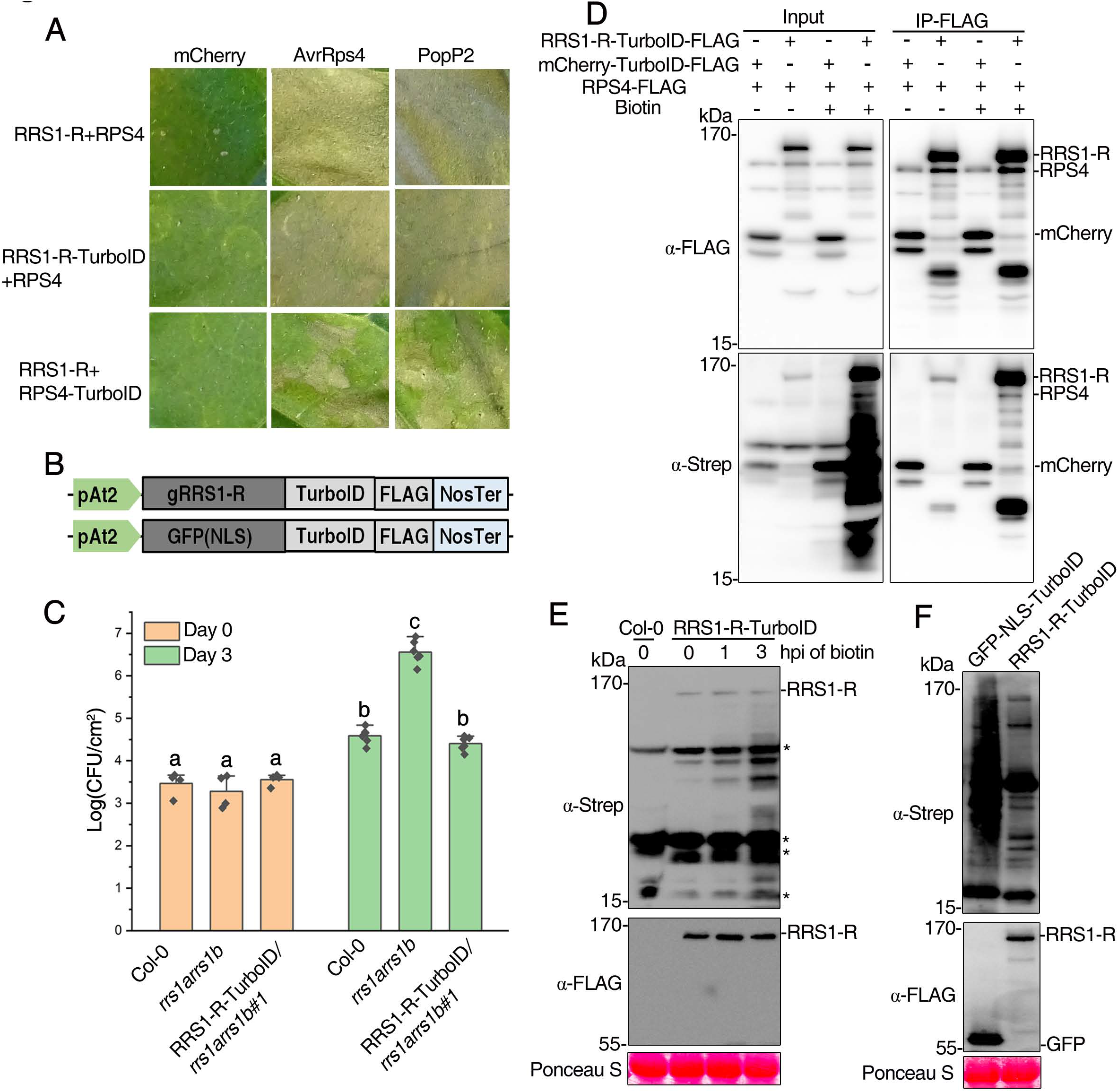
Establishment of TurboID-mediated proximity labeling in Arabidopsis for identification of proximal proteins of RRS1/RPS4 immune receptor complex. **(A)** Analysis of ability of RRS1-R-TurboID or RPS4-TurboID fusion construct to induce cell death in response to AvrRps4 and PopP2. Each tobacco leaf section was coinfiltrated to transiently express the indicated constructs combination together with either mCherry, AvrRps4 or PopP2. Photograph of the infiltrated leaf was taken at 4 days post infiltration (dpi). **(B)** Diagram of the expression cassettes used for the expression of TurboID biotin ligase. GFP protein containing a nuclear localization sequence (NLS) was fused to the N-terminus while a FLAG tag was added to the C-terminus of TurboID. Expression was under the control of Arabidopsis SSR16 (Small Subunit Ribosomal Protein 1 promoter, named as pAt2 in TSL ‘moderate promoter’ database for moderate expression control in transgenic Arabidopsis) and nopaline synthase terminator (NosTer) **(C)** RRS1-R-TurboID fusion protein complements loss of resistance to *Pst* DC3000 (AvrRps4) observed in the *rrs1arrs1b* mutant. Bacterial growth was measured 3 days post infiltration (dpi). Statistical significance is indicated by letters (p < 0.01, one-way ANOVA followed by Tukey’s post hoc test). **(D)** TurboID-based analysis of RPS4 biotinylation by RRS1-R-TurboID in *N. benthamiana*. *N. benthamiana* leaves were agroinfiltrated with the indicated constructs combination and biotin was infiltrated into the previously agroinfiltrated leaves at 36 h post-agroinfiltration (hpi). Agrobacteria containing mCherry-TurboID infiltrated into the leaves served as a negative control. IP was carried out using leaf samples collected 3 h after biotin treatment with anti-FLAG beads. The FLAG-tagged proteins were detected using anti-FLAG antibody and biotinylated proteins were detected using Streptavidin-HRP antibody, respectively. The experiment was repeated three times with similar results. **(E)** Biotinylation of RRS1-R and its vicinal proteins in RRS1-R-TurboID-FLAG transgenic plants. Total protein extracts from seedlings expressing RRS1-R–TurboID-FLAG with (+) or without (−) biotin treatment were immunoblotted with Streptavidin-HRP and anti-FLAG antibodies for detection of biotinylated proteins (top panel) and expression of TurboID (lower panel), respectively. Non-treated seedlings were used as controls (-) to visualize the background activity of the TurboID with endogenous biotin. Non-transformant (Col-0) seedlings were used as a control. The asterisks indicate the positions of naturally biotinylated proteins. **(F)** Streptavidin pull-down analysis of biotinylated proteins by TurboID-tagged RRS1-R. GFP-fused TurboID served as a control for the specificity of RRS1-R-TurboID-based proximity labeling. Western blot analysis of the Streptavidin pull-down products from leaf samples collected 3 h after biotin treatment probed with Streptavidin-HRP (top panel) and anti-FLAG antibody (lower panel). The molecular weight size markers in kDa are indicated at the left of each panel.

RPS4 and PopP2 are known to interact with RRS1^14,29^. To assess the specificity of TurboID-based proximity labeling, RPS4 or PopP2 were co-expressed with TurboID tagged RRS1-R in *N. benthamiana* followed by immunoprecipitation and biotinylation detection. Clear biotinylation signals were observed for RPS4 and PopP2 in the presence of TurboID tagged RRS1-R but not negative control mCherry (Figure 1D and S1B), indicating the high specificity of TurboID-based proximity labeling. Notably, consistent with our previous studies^14^, RRS1-R was not co-precipitated with PopP2, likely due to their weak or transient acetyltransferase–substrate interactions (Figure S1B). This finding supports the notion that proximity labeling outperforms affinity precipitation in identifying interactors with weak and transient interaction.

We then employed the *pAT2-RRS1-R-TurboID-FLAG* complementation in Col-0 *rrs1arrs1b* for the proximity-labeling proteomics. Wild-type plants expressing GFP carrying a nuclear localization signal (NLS) and TurboID-FLAG tag served as a negative control, as RRS1-R localizes in the nucleus^16^. In agreement with previous reports^30^, TurboID produced background labeling without the addition of exogenous biotin but the labelling yield can be further strongly increased in the presence of exogenous biotin. Biotin treatment of transgenic seedlings induced self-biotinylation of RRS1-R and robust whole-cell biotinylation (Figure 1E). The total biotinylated proteins were enriched using streptavidin-conjugated beads (Figure 1F) and analyzed via liquid chromatography/tandem mass spectrometry analysis. Proteins showing > two-fold higher peptide spectrum count in the RRS1-TurboID samples versus the GFP-TuboID samples were considered as RRS1-proximal proteins, leading to 284 candidates (Supplemental Table 1). Notably, among them are RPS4, a known interactor of RRS1-R, and regulators of plant NLRs, including Topless-related (TPR) proteins, MAC1 and MAC5A, suggesting the reliability of our data^31,32^. The balanced action of E3 ubiquitin ligases and deubiquitinases determines protein levels and activity^33^. A previously uncharacterized E3 ubiquitin ligase (AT1G18660, hereafter referred as RARE, <u>R</u>RS1-<u>a</u>ssociated <u>R</u>ING-type <u>E</u>3 ligase) and two redundant deubiquitinating enzymes (DUBs), ubiquitin-specific protease 12 (UBP12) and UBP13 are also among the candidates. Given their elusive nature on homeostasis regulation, we focused on further investigating RARE and UBP12/13.

### RARE is an E3 ubiquitin ligase that interacts directly with RRS1 and indirectly with RPS4 via RRS1

RARE contains a C3HC4-type RING domain with eight metal-binding residues (7 Cys and 1 His) coordinating two zinc ions in a cross-brace arrangement (Figure S2A), indicative of a potential E3 ubiquitin ligase^34^. A common feature of most E3 ligases is their ability to autoubiquitinate. *In vitro* ubiquitination assays revealed that RARE underwent autoubiquitination, while the RARE^H213Y^ variant (Figure 2A), harboring a tyrosine substitution of the zinc-binding histidine 213 (H213Y), did not (Figure S2A, Figure 2A). These results indicate that RARE is an active E3 ligase.

**Figure 2.**
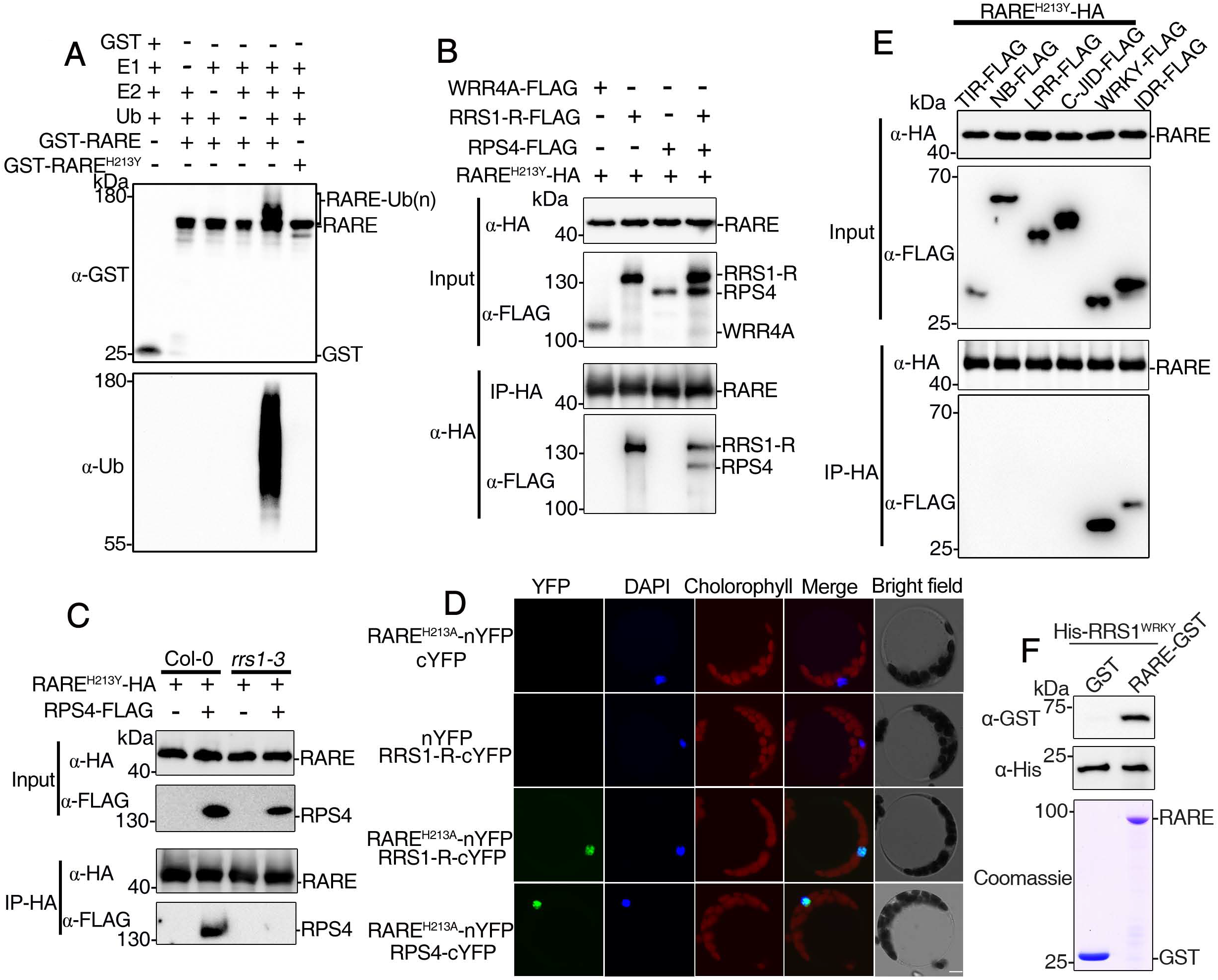
RARE is an E3 ubiquitin ligase that interacts directly with RRS1 and indirectly with RPS4 via RRS1. **(A)** Assays of in vitro self-ubiquitination of RARE. GST-RARE and its mutant form GST-RARE (H213Y) fusion proteins were assayed for E3 activity in the presence of E1 (UBA2), E2 (UBC10), ubiquitin (Ub) and ATP. “ + ” and “-” denote the presence or absence of the components of each reaction mixture. GST itself was used as a negative control. The presence of GST-RARE or its mutant form protein was detected using anti-GST antibody (upper panel). Protein ubiquitination bands generated by GST-RARE are indicated on the right. Ubiquitination results in a heterogeneous collection of higher molecular mass proteins that are detected with anti-ubiquitin antibody (lower panel). **(B)** Co-immunoprecipitation(co-IP) assays reveals that the interaction of RARE with RRS1/RPS4 complex is dependent on RRS1. Samples were harvested from *Nb* leaves transiently coexpressing FLAG-tagged RRS1/RPS4 and HA-tagged RARE. Total extracts were immunoprecipitated with agarose-conjugated anti-HA antibody followed by immunoblotting with the indicated antibodies. WRR4A was used as a control. **(C)** Co-IP assays for the interaction of RARE with RPS4 in protoplasts isolated from Col-0 and *rrs1-3* plants. Protoplasts isolated from Col-0 and *rrs1*-3 co-transfected with the RARE-HA and RPS4-FLAG constructs were incubated overnight, and total protein extracts were subjected to immunoprecipitation with agarose-conjugated anti-HA antibody followed by immunoblot analysis using either anti-HA or anti-FLAG antibodies. **(D)** BiFC analyses for the interaction of RARE with RRS1/RPS4 complex in Arabidopsis Col-0 protoplasts. Arabidopsis mesophyll protoplasts were transformed with the indicated BiFC constructs and the YFP fluorescence was visualized by confocal microscopy 16-20 h after transient expression. The positions of nuclei were shown by 4, 6-diamidino-2-phenylindole (DAPI) staining. Scale bar, 5 μm. **(E)** Co-IP assays to evaluate the association of HA-tagged RARE with FLAG-tagged individual domain of RRS1-R after transient co-expression in *Nb* leaves. Total protein extracts were subjected to immunoprecipitation with agarose-conjugated anti-HA antibody followed by immunoblot analysis using either anti-HA or anti-FLAG antibodies. **(F)** Pull-down assay for the interaction of RARE with WRKY domain of RRS1. His-tagged WRKY was incubated with immobilized GST or GST-tagged RARE. After washing, bound proteins were eluted and subjected to immunoblot analysis using an antibody against His. Coomassie blue (CBB) staining indicates equal amounts of bait proteins

The identification of RARE from proximity labeling proteomics prompted us to test its interaction with RRS1-R and RPS4. We performed co-immunoprecipitation (Co-IP) assays after transient coexpression of E3 ligase activity-deficient RARE^H213Y^ with RRS1-R or RPS4 in *N. benthamiana* since a functional E3 ligase might destabilize its substrate and reduce protein–protein interactions strength^35^. RARE^H213Y^ interacted with RRS1-R but not with RPS4 and the negative control TNL protein WRR4A (White Rust Resistance 4A)^36^, However, the interaction between RARE^H213Y^ and RPS4 could only be observed in the presence of RRS1-R (Figure 2B), suggesting that RARE interacts with RPS4 via RRS1-R. The RRS1-dependent RARE-RPS4 interaction was further supported by finding that RPS4 associated with RARE^H213Y^ in Co-IPs conducted in Arabidopsis protoplast derived from wild-type plants but not from the *rrs1-3* mutants that lack the endogenous RRS1 proteins ^28^(Figure 2C). Consistently, the fluorescence complementation (BiFC) assays showed that RARE^H213Y^ interacts with RPS4 in protoplasts derived from wild-type plants but not from the *rrs1-3* mutants, while its association with RRS1-R was observed in nucleus (Figure 2D and S2B). This agrees with the nuclear localization of RRS1-R and the nucleocytoplasmic localization of RARE (Figure S2C and S2D).

To identify the domain of RRS1-R responsible for RARE interaction, we performed Co-IPs in *N. benthamiana* by co-expressing RARE^H213Y^ with individual domains of RRS1-R. In the Co-IP assays, RARE^H213Y^ interacts strongly with the WRKY domain and weakly with intrinsically disordered region (IDR), but not with other domains of RRS1-R (Figure 2E). As IDRs are prone to forming weak promiscuous interactions due to their high flexibility^37^, we conclude that RARE interacts with RRS1-R primarily through the WRKY domain. *In vitro* pull-down assays showed that the WRKY of RRS1-R directly binds to RARE^H213Y^ (Figure 2F), indicative of a direct interaction between RARE and RRS1-R. Collectively, these data suggest that RARE interacts directly with RRS1-R primarily through its RRS1-R^WRKY^, and indirectly with RPS4 via RRS1-R.

### RARE ubiquitinates RRS1 through its integrated WRKY domain

The direct interaction between RARE and RRS1 led us to examine whether RARE ubiquitinates RRS1. When co-expressed in *N. benthamiana*, RRS1-R was ubiquitinated by RARE but not the E3 ligase-dead RARE^H213Y^ variant (Figure 3A), suggesting that RRS1-R is ubiquitinated by RARE. To confirm this in Arabidopsis, we obtained the *rare* mutant (a T-DNA null mutant of *RARE*, Figure S3A and B), and crossed it into wild-type plants expressing *pAT2::RRS1-R-FLAG* generated by our previous study^18^. Ubiquitinated RRS1-R was detected in the *pAT2::RRS1-R-FLAG* transgenic lines, while the introduction of *RARE* knockout almost abolished RRS1-R ubiquitination (Figure 3B). Collectively, these results indicate that RARE ubiquitinates RRS1-R.

**Figure 3.**
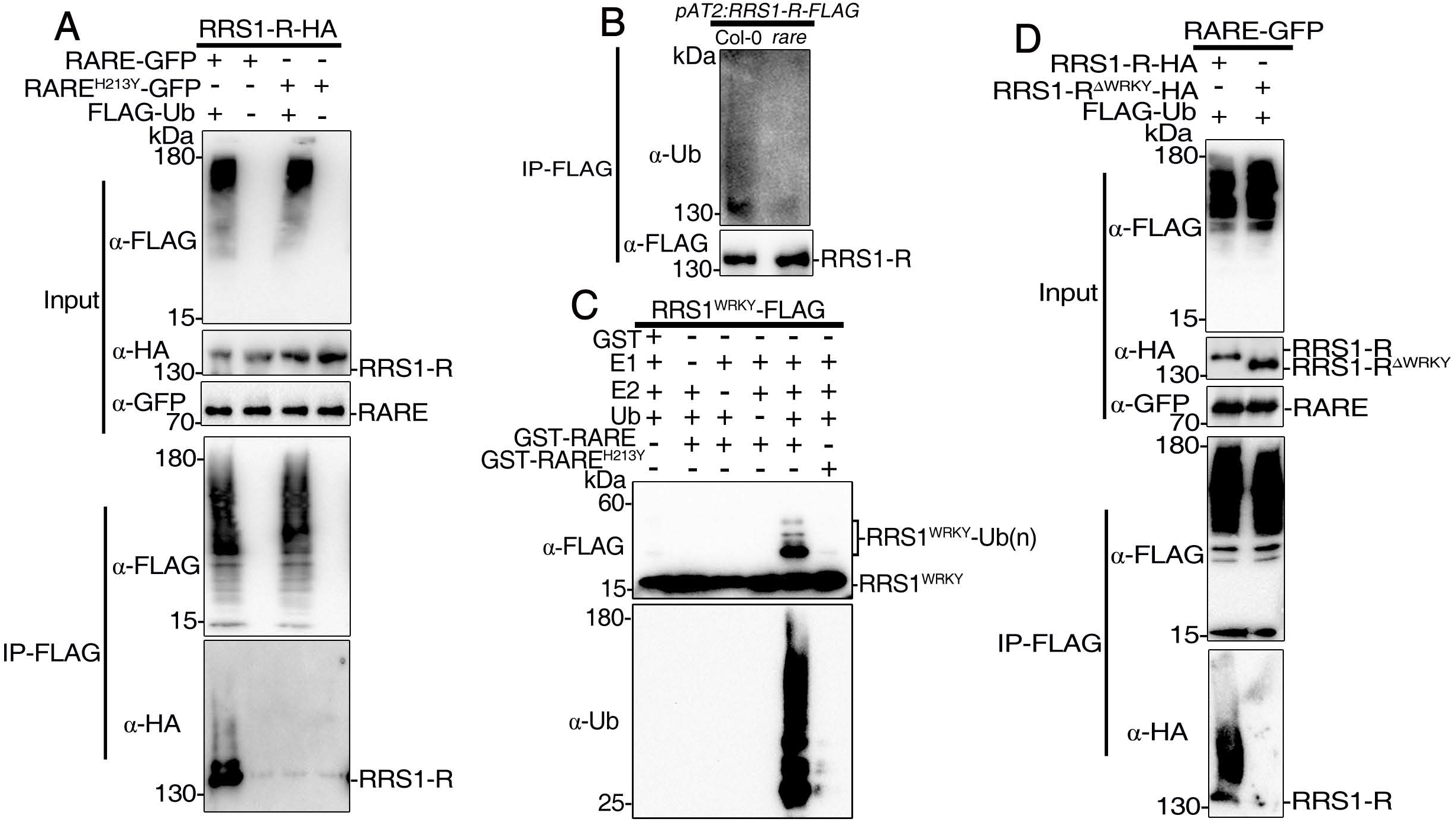
RARE is a functional E3 ligase and ubiquitinates the WRKY domain of RRS1. **(A)** Detection of ubiquitination of RRS1-R by RARE in *Nb* leaves. HA-tagged RRS1-R was co-expressed with GFP-tagged RARE or its mutant form RARE (H213Y) in the presence or absence of FLAG-Ub in *Nb* leaves. The *Nb* leaves were pretreated with 50 μM MG132 for 6h before harvesting. Total ubiquitinated proteins were immunoprecipitated at 36h post-infiltration with anti-FLAG antibody, and ubiquitinated RRS1-R proteins were detected by immunoblotting with anti-HA antibody. **(B)** Reduced *in vivo* ubiquitination level of RRS1 in *rare* mutant compared to wild-type background. Total protein extracts from FLAG-tagged RRS1 plants in wild-type or *rare* background were subjected to immunoprecipitation using anti-FLAG antibody. Following immunoprecipitation with anti-FLAG antibody, the ubiquitination of RRS1 was detected by immunoblot analysis using anti-Ubiquitin antibody. Immunoblots (IB) were probed with anti-ubiquitin and anti-FLAG antibody, respectively. **(C)** Ubiquitination of WRKY domain of RRS1 by RARE *in vitro*. FLAG-tagged WRKY domain of RRS1 was incubated with GST-RARE in ubiquitination assay buffer. Samples were resolved by SDS-PAGE and subjected to immunoblot analysis with anti-FLAG (top panel) or anti-Ub (bottom panel) antibody. Direct ubiquitination of WRKY domain was evident by higher molecular laddering detected by immunoblotting with anti-FLAG antibody. **(D)** WRKY domain is required for RRS1 ubiquitination by RARE. HA-tagged RRS1-R or its WRKY domain deletion variant was co-expressed with GFP-tagged RARE in the presence of FLAG-Ub in *Nb* leaves. The *Nb* leaves were pretreated with 50 μM MG132 for 6h before harvesting. Total ubiquitinated proteins were immunoprecipitated at 36h post-infiltration with anti-FLAG antibody, and ubiquitinated RRS1-R proteins were detected by immunoblotting with anti-HA antibody.

Since the interaction with RARE is primarily mediated by the integrated WRKY domain of RRS1-R, we next tested whether RARE could directly ubiquitinate RRS1-R^WRKY^. *In vitro* ubiquitination assays showed that RRS1-R^WRKY^ was ubiquitinated by RARE but not by RARE^H213Y^ (Figure 3C). Additionally, RARE ubiquitinated RRS1-R but not the RRS1-R^ΔWRKY^ variant lacking the WRKY ID when co-expressed in *Nb* (Figure 3D). These data suggest that RARE ubiquitinates RRS1-R through its WRKY domain.

Many Arabidopsis accessions carry RRS1B/RPS4B, an RRS1/RPS4 paralogous pair with similar domain architecture. The WRKY domains of RRS1B and RRS1 share ∼ 60% amino acid sequence identity^28^. Like RRS1-R, we found that both RRS1B and RRS1B^WRKY^ can be ubiquitinated by RARE (Figure S3C and S3D), albeit RRS1B^WRKY^ ubiquitination is less efficient than that of RRS1-R^WRKY^ (Figure S3E). These data suggest that RARE ubiquitinates RRS1 and its paralogue RRS1B through their integrated WRKY domains.

### RARE compromises the abundance and immune responses of the RRS1/RPS4 complex

E3 ligase-catalyzed ubiquitination often targets substrates for proteasomal degradation^38^. As RARE ubiquitinates RRS1-R, we tested its effect on RRS1/RPS4 complex abundance. When co-expressed in *Nb*, accumulation of RRS1-R was impaired by EARE. Intriguingly, RARE destabilized RPS4 only when co-expressed with RRS1-R, but not RPS4 alone (Figure 4A). This destabilization requires RARE’s ubiquitin ligase activity, as RARE^H213Y^ failed to reduce the complex levels (Fig S4A). Consistent with our previous studies^39,40^, we found that RRS1 stabilized RPS4 (Figure 4A). RARE likely destabilizes RPS4 by promoting RRS1-R degradation, thereby disrupting RRS1-R-mediated stabilization of RPS4. Supporting this, transiently expressing RARE reduced the protein abundance of RPS4 in wild-type but not *rrs1-3* Arabidopsis protoplasts (Figure 4B). Collectively, these results demonstrate that RARE negatively regulates RRS1/RPS4 complex abundance in an RRS1-dependent manner.

**Figure 4.**
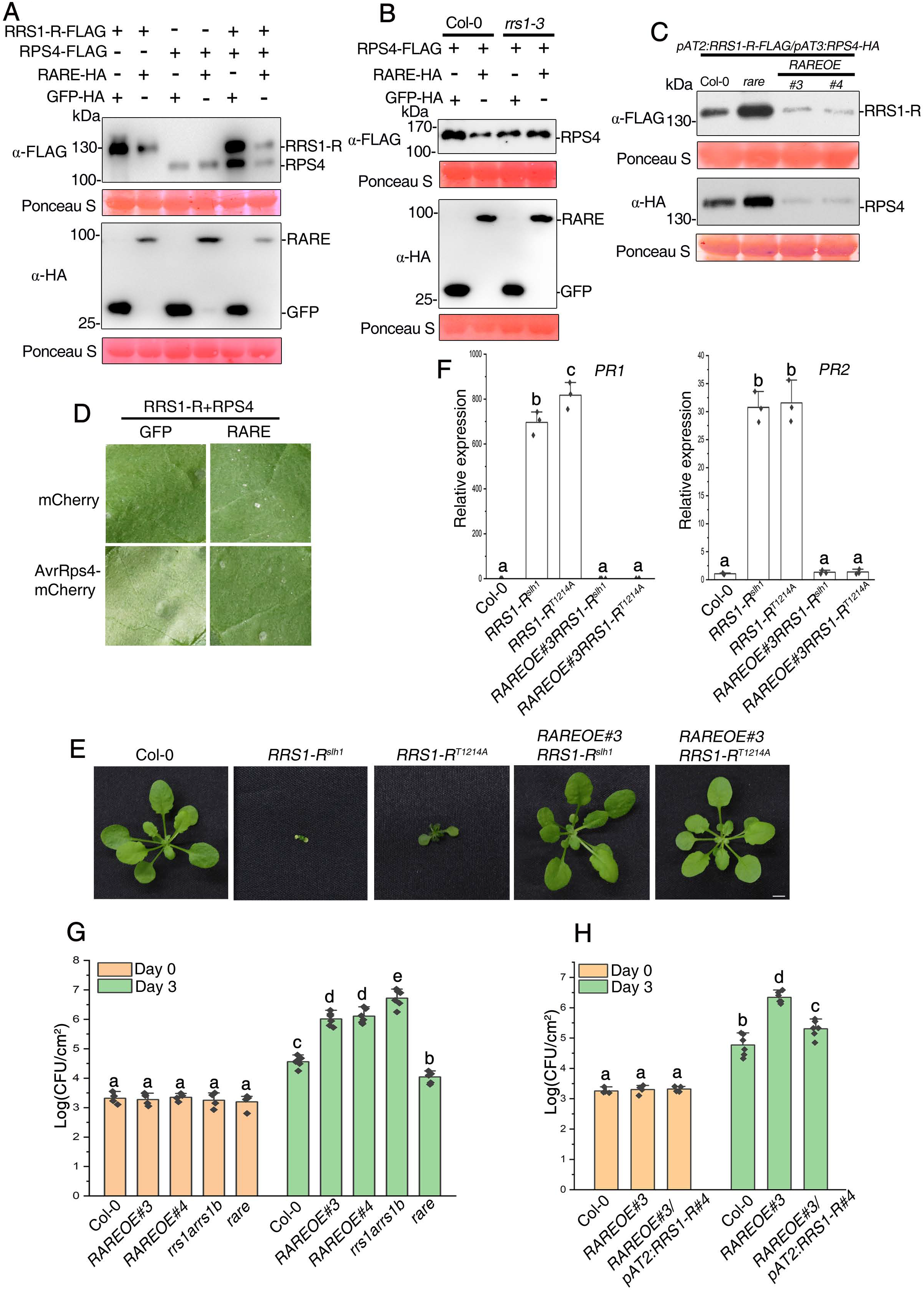
RARE destabilizes RRS1/RPS4 complex in a RRS1-dependent manner and negatively regulates RRS1/RPS4–mediated defense responses. **(A)** RARE directly destabilizes RRS1 and indirectly destabilizes RPS4 in the presence of RRS1. RRS1-R, RPS4 or RRS1-R/RPS4 were transiently co-expressed with RARE in *Nb* at 2 dpi and then subjected to immunoblotting with anti-FLAG antibody for detecting RRS1-R/RPS4 or anti-HA antibody for detecting RARE. The HA-tagged GFP served as a negative control. **(B)** RARE modulates RPS4 homeostasis via RRS1. RPS4-FLAG was co-expressed together with RARE-HA or GFP-HA in protoplasts isolated from Col-0 or *rrs-1* plants. RPS4-FLAG proteins were detected by immunoblotting with anti-FLAG antibody. **(C)** Immunoblot analyses of RRS1-R and RPS4 protein levels in Col-0, *rare*, and *RAREOE* background. Total proteins were extracted from 10-days-old seedlings and subjected to immunoblot analysis using anti-FLAG antibody for detecting RRS1-R and anti-HA antibody for detecting RPS4, respectively. Ponceau S staining of Rubisco indicates equal loading. **(D)** Suppression of RRS1/RPS4-mediated cell death upon AvrRps4 perception in tobacco plants. Each leaf section was transiently coinfiltrated with RRS1-R/RPS4, AvrRps4, and either GFP or RARE. Photographs assessing HR were taken 4 dpi. Similar results were obtained in two additional experiments. **(E)** Overexpression of RARE suppresses the autoimmune phenotype triggered by RRS1-R^slh1^ and RRS1-R^T1214A^, the auto-active alleles of RRS1-R. Morphology of 5-week-old soil-grown plants of wild-type (WT), *RRS1-R^slh1^*, *RRS1-R^T1214A^*, *RAREOERRS1-R^slh1^* and *RAREOERRS1-R^T1214A^*. Scale bar, 0.5 cm. **(F)** Relative expression levels of *PR1* and *PR2* were determined by real-time PCR. The relative transcript level of *PR1* and *PR2* were normalized to the expression of *UBQ*. Error bars represent SD from three measurements. The gene expression analysis was performed on three batches of independent grown plants. **(G)** *RAREOE* lines are compromised in RRS1/RPS4-mediated resistance against *Pst* DC3000(AvrRps4). 5-week-old plants were infiltrated with the bacteria at OD_600_= 0.0005. Leaf discs within the infiltrated area were taken at days 0 and 3 to measure the bacterial growth. Bars represent the SD of six replicates. **(H)** Introduction of the *RRS1-R* transgene into the *RAREOE3* line restores disease resistance to *Pst* DC3000(AvrRps4). Note that the epitope-tagged RRS1-R/RPS4 transgenic line#4 was used to cross with *rare* and *RAREOE* transgenic line.

To verify RARE’s homeostatic control of the RRS1/RPS4 complex in Arabidopsis, we generated *RARE*-overexpressing transgenic lines driven by the constitutive 35S promoter (*RAREOE*, Fig S4B and S4C) and crossed the *RAREOE* transgene and the *rare* null allele into the epitope-tagged RRS1-R/RPS4 transgenic lines (*pAT2::RRS1-R-FLAG/pAT3::RPS4-HA*) generated by a previous study^18^. RRS1 and RPS4 abundance considerably increased in *rare* but decreased in *RAREOE* background (Figure 4C), while *RRS1-R* transcripts remained similar (Fig S4D), suggesting that RARE post-translationally modulates RRS1/RPS4 complex levels. Cycloheximide (CHX) chase assays revealed slower degradation of RRS1-R/RPS4 complex in *rare* than in wild-type background and the degradation was inhibited by the 26S proteasome inhibitor MG132 (Figure S4E-F). Taken together, these results demonstrate that RARE regulates RRS1/RPS4 abundance by mediating RRS1 degradation via the 26S proteasome machinery.

RARE’s homeostatic control of the RRS1/RPS4 complex prompted us to test its role in regulating RRS1/RPS4–mediated defense responses. Tobacco transient expression showed that RARE abolished HR triggered by RRS1/RPS4 in recognition of AvrRps4 (Figure 4 D). RRS1-R^slh^^1^ and RRS1-R^T1214A^ are two auto-active alleles of RRS1-R that carry mutations in the WRKY domain^18,41^, thus activating the RRS1-R/RPS4 complex when transgenically expressed in Arabidopsis Col-0 plants. Crossing *RAREOE* into the transgenic plants expressing RRS1-R^slh^^1^ and RRS1-R^T1214A^ rescued their autoimmune phenotypes, including dwarfism and elevated *Pathogenesis-related Proteins 1* (*PR1*) and *PR2* expression (Figure 4E-F). The endogenous RRS1/RPS4 complex confers resistance against bacterial pathogen *Pst* DC3000(AvrRps4) in wild-type plants. Compared to wild-type plants, the *RAREOE* plants exhibited compromised resistance, while the *rare* mutants displayed enhanced resistance (Figure 4G). Taken together, these data suggest RARE negatively regulates HR and resistance-mediated by the RRS1/RPS4 complex (Figure S4F).

To verify that the diminished RRS1 accumulation accounts for impaired RRS1/RPS4-mediated immunity in the *RAREOE* plants, we crossed the *RAREOE* lines with FLAG-tagged *RRS1-R* transgenic line #4 with higher RRS1-R protein levels (Figure S4G). This largely restored resistance to *Pst* DC3000 carrying AvrRps4 in the *RAREOE* plants(Figure 4H), indicating that the susceptibility of the *RAREOE* lines was caused by diminished RRS1 accumulation. Taken together, our data indicate that RARE destabilizes the RRS1/RPS4 complex by promoting RRS1 degradation, thus compromising RRS1/RPS4-mediated defense responses.

### UBP12/UBP13 counteract RARE’s regulation of RRS1/RPS4

Deubiquitinating enzymes (DUBs) remove ubiquitin from ubiquitinated substrates, reversing the ubiquitination process. The Arabidopsis ubiquitin-specific proteases (UBPs) belong to the largest subfamily of DUBs and regulate diverse cellular processes^42,43^. Like E3 ligase RARE, UBP12 and UBP13, two functionally redundant UBPs^44^, were identified in the proximal proteome of RRS1-R (Supplemental Table 1). We therefore tested if they counteract RARE’s regulation of RRS1/RPS4. Semi-*in vitro* deubiquitination assays revealed that immunoprecipitated RRS1 from the *pAT2::RRS1-R-FLAG* plants was deubiquitinated by UBP12 and UBP13, but not their deubiquitinase-dead variants UBP12^C208S^ and UBP13^C207S^(Figure 5A). Neither UBP12 nor UBP13 affected autoubiquitination of RARE (Figure S5A). These data indicates that UBP12 and UBP13 directly deubiquitinate RRS1 in a deubiquitinase activity-dependent manner without affecting RARE’s activity. RARE ubiquitinates RRS1 on its integrated WRKY domain. We next examined if UBP12/UBP13 can remove RARE-catalyzed ubiquitination of RRS1-R^WRKY^. The ubiquitinated RRS1-R^WRKY^, generated through *in vitro* ubiquitination reaction with RARE and subsequently purified, was dramatically reduced by UBP12 or UBP13, but not their deubiquitinase-dead variants nor the negative control UBP7 (Figure 5B, Figure S5B), as revealed by the *in vitro* deubiquitination assays. These results demonstrate that UBP12/UBP13 antagonize RARE-mediated ubiquitination of RRS1-R^WRKY^.

**Figure 5.**
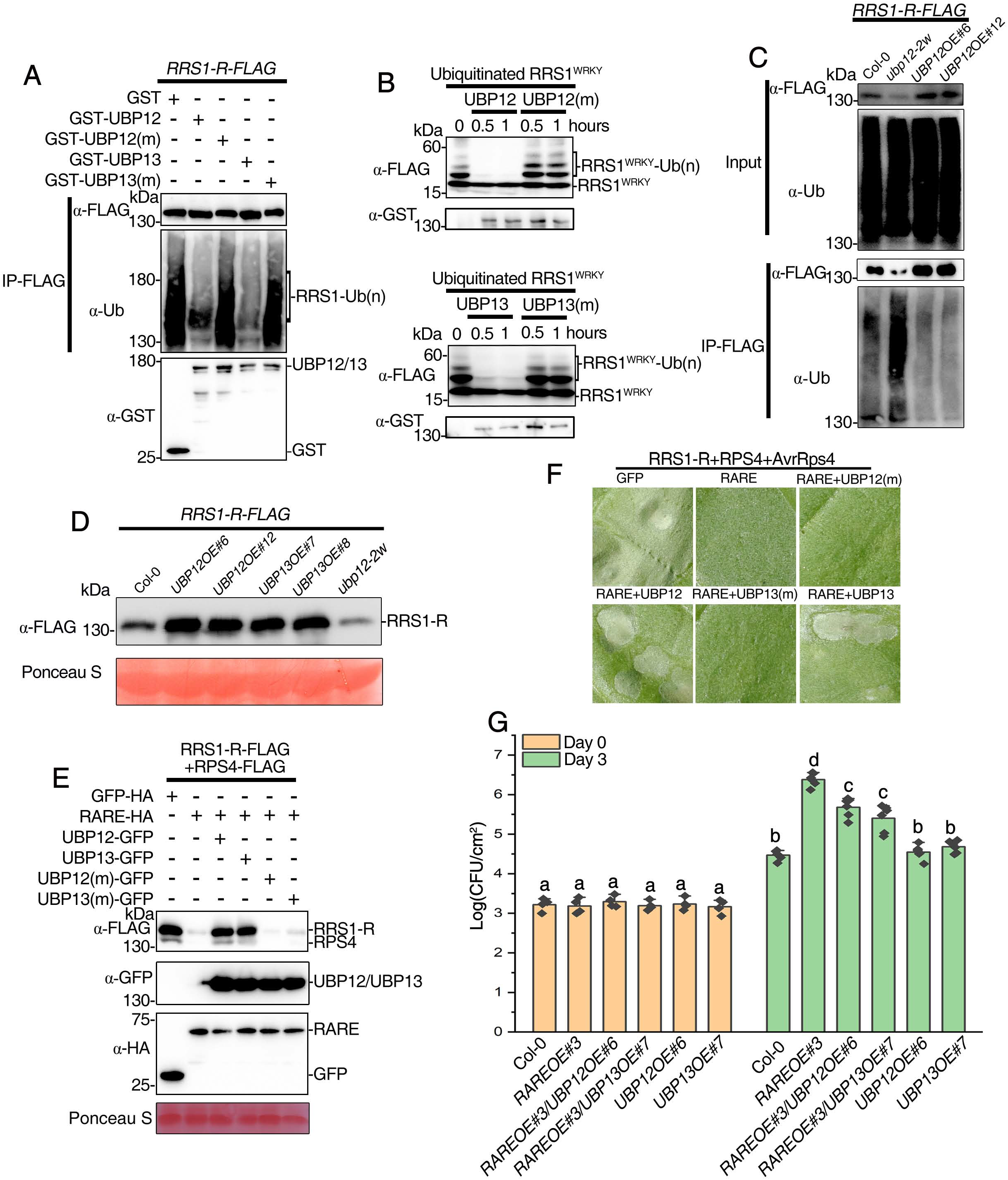
UBP12 and UBP13 counteract the effect of RARE by removing ubiquitin from polyquibiquinated RRS1. **(A)** Deubiquitination of RRS1 expressed in Arabidopsis plants by recombinant UBP12 and UBP13. Polyubiquitinated RRS1 proteins were immunoprecipitated with agarose-conjugated anti-FLAG antibody from 10-day-old seedlings and subsequently co-incubated with GST-UBP12/UBP13 or GST-UBP12C208S/UBP13C207S for 2 h. Immunoprecipitated RRS1 was detected by anti-FLAG antibody (top panel). RRS1 ubiquitination status was analyzed by immunoblotting with anti-ubiquitin antibody (middle panel). The presence of GST-tagged UBP recombinant proteins was confirmed by the α-GST antibody (bottom panel). **(B)** UBP12 and UBP13 deubiquitinate polyubiquitinated WRKY domain of RRS1 in vitro. Polyubiquitinated WRKY domain proteins produced by ubiquitination in the presence of E1(UBA2), E2 (UBC10), RARE and ubiquitin (Ub) were pulled down and then incubated with GST-UBP12, GST-UBP13, or their catalytically inactive variants for 0.5 or 1 hour. Ubiquitinated WRKY domain was detected by anti-FLAG antibody (top panel). GST-UBP12, GST-UBP13 or the inactive variants were detected with anti-GST antibody (bottom panel). **(C)** The polyubiquitination status of RRS1 in various genotypes. 10-days-old seedlings grown on MS medium were pretreated with 50 mM MG132 for 6 h. Protein extracts were immunoprecipitated with agarose-conjugated anti-FLAG antibody, followed by immunoblotting with anti-ubiquitin (top panel) and anti-FLAG antibodies (bottom panel). **(D)** The protein levels of FLAG-tagged RRS1 driven by pAT2 promoter in various genotypes. Total protein extracts from 10-days-old seedlings were subjected to immunoblotting with anti-FLAG antibody for detecting RRS1-R. Two independent lines were used in (C) and (D) separately. **(E)** UBP12/UBP13 antagonize RARE-mediated RRS1/RPS4 complex degradation. RRS1/RPS4 was co-expressed with RARE in *Nb* leaves in the presence or absence of UBP12/UBP13. GFP was coexpressed in the experiment as a control. **(F)** UBP12/UBP13 attenuate RARE-mediated suppression of cell death caused by coexpression of RRS1-R/RPS4 and AvrRps4. Each leaf section was transiently coinfiltrated with RRS1-R/RPS4, AvrRps4, and the indicated constructs. Photographs assessing HR were taken 4 dpi. **(G)** Bacterial growth of *Pst* DC3000(AvrRps4) on 5-week-old leaves of the indicated genotypes at 0 and 3 dpi with bacterial inoculum of OD_600_=0.0005. Statistical significance is indicated by different letters (P < 0.01, one-way ANOVA followed by Tukey’s post hoc test). Bars represent means ± SD. All the experiments were repeated three times with similar results.

To investigate the deubiquitination of RRS1 by UBP12/UBP13 *in vivo*, the *pAT2::RRS1-R-FLAG* transgene was crossed into *UBP12OE*, *UBP13OE* transgene lines and *ubp12-2w*, a weak double mutant with reduced *UBP12* and *UBP13* transcripts, since the *ubp12ubp13* double null mutants are infertile^45^. Polyubiquitinated RRS1-R levels were greatly increased in *ubp12-2w* and reduced in *UBP12OE* and *UBP13OE* compared to those in wild-type (Figure 5C and S5C). To correlate UBP12/UBP13 with RRS1 abundance, we compared RRS1 protein abundance in these backgrounds. RRS1 accumulation was significantly increased in *UBP12O*E and *UBP13OE* but reduced in *ubp12-2w* (Figure 5D). The addition of MG132 partially restored RRS1 abundance in *ubp12-2w* (Figure S5D). Consistently, UBP12 and UPB13 rescued the RARE-mediated reduction of RRS1-R and RPS4 in a deubiquitinase activity-dependent manner (Figure 5E). Collectively, these results suggest that UBP12 and UBP13 deubiquitinate RRS1, protecting it from proteasomal degradation and thereby stabilizing the RRS1/RPS4 complex.

Finally, we tested whether UBP12/UBP13 affects RRS1/RPS4-mediated immune responses. HR assay in tobacco showed that UBP12/UBP13, but not their mutants, alleviated RARE-mediated suppression of HR triggered by RRS1-R/RPS4 in recognition of AvrRps4 (Figure 5F). Additionally, introducing *UBP12OE* and *UBP13OE* into the *RAREOE* lines by crossing partially rescued the enhanced susceptibility of the *RAREOE* lines against *Pst* DC3000(AvrRps4) (Figure 5G). These results indicate that UPB12/13 counteract RARE’s inhibition of RRS1/RPS4-mediated immune responses. Taken together, our data suggests that UBP12 and UBP13 antagonize RARE-mediated ubiquitination of RRS1-R, consequently protecting the accumulation and defense responses of the RRS1/RPS4 complex.

### RARE and UBP12/UBP13 antagonistically regulate two WRKY transcription factors homologous to RRS1-R^WRKY^

IDs, usually homologous to effector target proteins, were incorporated into NLR-IDs through domain shuffling for effector detection during evolution^23^. Phylogenetic analysis revealed that RARE and UBP12/UBP13 are both conserved across monocots and dicots (Fig S6A and S6B). In contrast, RRS1 homologs carrying the integrated WRKY domain are limited to the Camelineae tribe. Outside of this tribe, RRS1 homologs lack the WRKY ID (Figure 6A), suggesting that WRKY domain integration likely occurred during the diversification of Camelineae tribe within the Brassicaceae family. RARE and UBP12/UBP13 antagonistically regulate the reversible ubiquitination of RRS1 through ubiquitinating/deubiquitinating the WRKY ID. Given the limited co-existence of RARE and UBP12/UBP13 with RRS1-R^WRKY^ domain during evolution, the initial roles of RARE and UBP12/UBP13 may be involved in homeostatic regulation effector targets homologous to RRS1-R^WRKY^.

**Figure 6.**
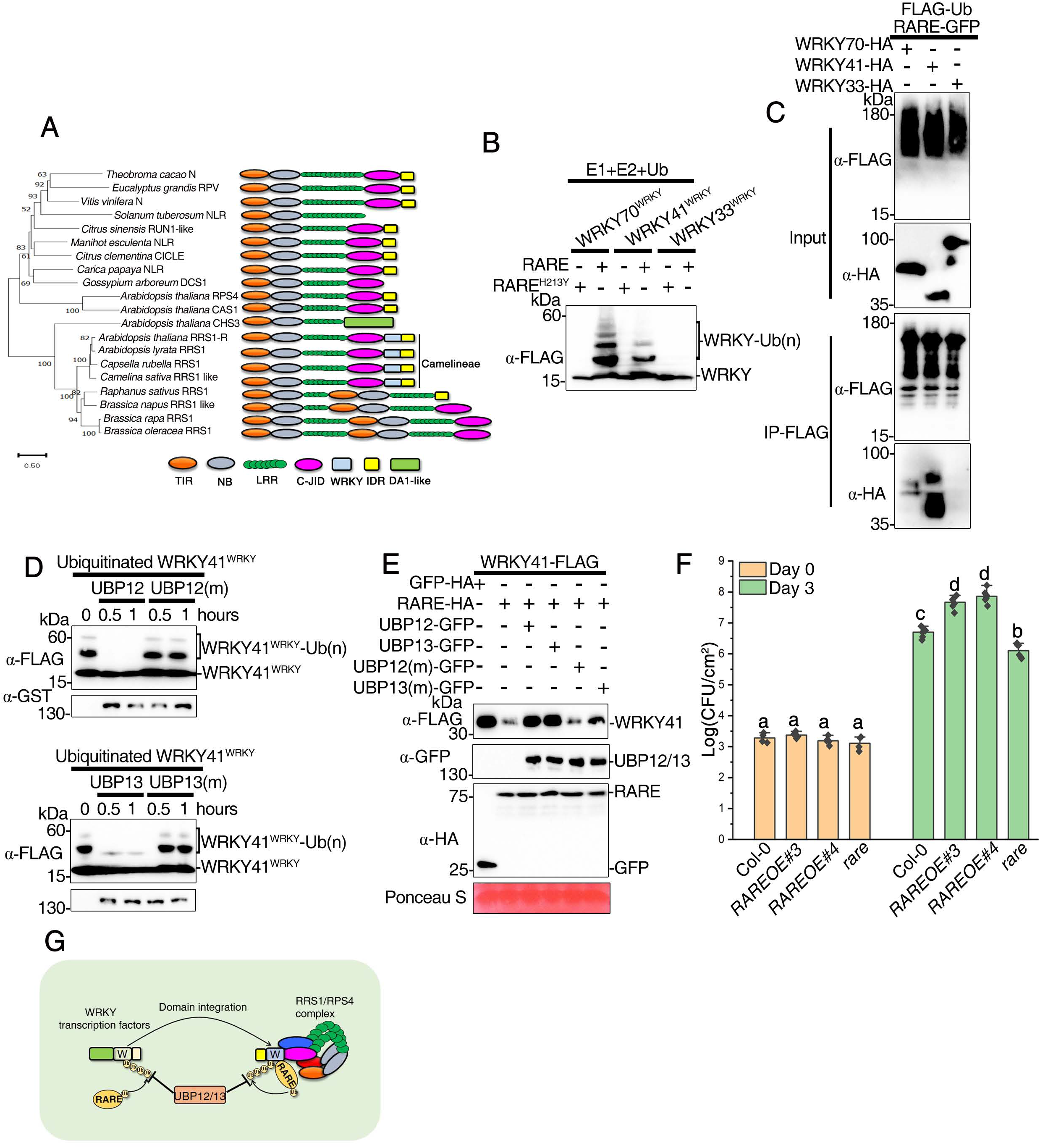
RARE and UBP12/UBP13 antagonistically regulate the stability some WRKY transcription factors homologous to RRS1-R^WRKY^. **(A)** The phylogenetic tree of RRS1 orthologues (left) and their domain architecture (right). The tree was generated using the neighbour-joining method based on full-length amino acid sequences. Only bootstrap values >50% within 1000 replicates are shown above branches. RPS4, CSA1 and CHS3 serve as outgoup. **(B)** RARE ubiquitinates the WRKY domain of WRKY70, WRKY41 but not WRKY33. FLAG-tagged WRKY domains were incubated with GST-RARE or its mutant form RARE (H213Y) in ubiquitination assay buffer containing E1(UBA2), E2 (UBC10) and Ub. Samples were resolved by SDS-PAGE and subjected to immunoblot analysis with anti-FLAG antibody for detecting higher molecular weight laddering bands. **(C)** Detection of ubiquitination of WRKY70 and WRKY41 but not WRKY33 by RARE in *Nb* leaves. HA-tagged WRKY proteins were co-expressed with GFP-tagged RARE in the presence of FLAG-Ub in *Nb* leaves. The *Nb* leaves were pretreated with 50 μM MG132 for 6h before harvesting. Total ubiquitinated proteins were immunoprecipitated at 36h post-infiltration with anti-FLAG antibody, and ubiquitinated WRKY proteins were detected by immunoblotting with anti-HA antibody. **(D)** UBP12 and UBP13 deubiquitinate polyubiquitinated WRKY domain of WRKY41 in vitro. Anti-FLAG and anti-Ub antibodies were used to detect the ubiquitination of WRKY domain. Polyubiquitinated WRKY domain proteins produced by ubiquitination in the presence of E1(UBA2),E2 (UBC10), RARE and ubiquitin (Ub) were pulled down and then incubated with GST-UBP12, GST-UBP13, or their catalytically inactive variants for 0.5 or 1 hour. Ubiquitinated WRKY domain was detected by anti-FLAG (top panel). GST-UBP12, GST-UBP13 or the inactive variants were detected with anti-GST antibody (bottom panel). **(E)** UBP12/UBP13 antagonize RARE-mediated WRKY41 degradation. WRKY41 was co-expressed with RARE in *Nb* leaves in the presence or absence of UBP12/UBP13. GFP was coexpressed in the experiment as a control. **(F)** RARE contributes to basal defense against virulent pathogens. Five-week-old leaves of the indicated genotypes were infiltrated with *Pst* DC3000 at a final concentration of OD_600_=0.0002. Leaf discs within the infiltrated area were taken at days 0 and 3 to measure the bacterial growth. Statistical significance is indicated by different letters (P < 0.01, one-way ANOVA followed by Tukey’s post hoc test). Bars represent means ± SD. **(G)** A model illustrating UBP12/UBP13- and RARE-mediated (de)ubiquitinating the WRKY ID of RRS1 as an evolutionarily gained modification for fine-tuning homeostasis of RRS1/RPS4 complex.

Our previous studies show that WRKY70, WRKY41 and WRKY33 are three putative targets of AvrRps4 and PopP2^16^. The WRKY domains of WRKY70 and WRKY41, but not WRKY33, are in the same clade as RRS1-R^WRKY^ in the phylogenetic tree (Figure S6C), suggesting that they are homologous to RRS1-R^WRKY^. Consistent with our speculation, ubiquitination assays showed that RARE is capable of ubiquitinating the WRKY domains and full-length of WRKY70, WRKY41 but not WRKY33 (Figure 6B, C). As expected, UBP12 and UBP13 also can remove ubiquitin from ubiquitinated WRKY domains of WRKY41 and WRKY70 catalyzed by RARE (Figure 6D, Figure S6D). These data suggest that WRKY70 and WRKY41 are under reversible ubiquitination regulation by RARE and UBP12/UBP13. Consistent with UBP12/UBP13 regulating the ubiquitination of WRKY41 and WRKY70 in a manner opposite to RARE, RARE promoted the degradation of WRKY70 and WRKY41, but not WRKY33, while UBP12/13 inhibited such degradation (Figure 6E, Figure S6E and S6F), indicating antagonistic control of WRKY70 and WRKY41 stability.

WRKY70 and WRKY41 positively regulate basal resistance^46,47^. Accordingly, the *rare* mutants exhibit enhanced basal resistance against *Pst* DC3000, while the *RAREOE* lines show reduced resistance (Figure 6F). Silencing of UBP12/UBP13 enhanced resistance against *Pst* DC3000^44^, which seems contradictory to the stabilization of WRKY70 and WRKY41 by UBP12/UBP13. The paradox likely arises from the compound effects of UBP12/UBP13 substrates functioning as negative regulators of basal resistance, including MYC2, NPR3 and NRP3, that are also stabilized by UBP12/UBP13^42,48^. Collectively, these results suggest that RARE and UBP12/UBP13 antagonistically regulate ubiquitination and stability of WRKY70 and WRKY41, which carry homologous WRKY domains to RRS1, and that such regulatory mechanism was co-opted to modulate RRS1/RPS4 complex homeostasis during WRKY domain integration into RRS1(Figure 6G). Therefore, WRKY ID-mediated ubiquitination of RRS1 has been retained in evolution and continues to contribute for fine-tuning homeostasis of RRS1/RPS4 complex

## Discussion

NLR homeostasis must be tightly regulated to ensure proper defense without triggering autoimmunity. Many NLRs have been shown to associate with diverse group of NLR-interacting proteins to achieve initiation and fine-tuning of immune responses, which forms another layer of regulation of NLR activity. In this study, we exploited TurboID-based proximity labeling proteomics to profile the proxiomes of RRS1/RPS4 complex using RRS1-R-TurboID transgenic plants. A number of new regulators potentially involved in the RRS1/RPS4-mediated immune response were identified. Some proteins that have been previously reported to be associated with other NLRs or to participate in the immune signaling, such as TPR proteins, MAC1 and MAC5A, were also identified in our PL-MS dataset^31,32^. So far, no interacting partner has been identified to modulate the activity or stability of RRS1/RPS4 complex. Therefore, the proteomic datasets described in this work provide an overview of regulatory proteins and signaling partners associated with RPS4/RRS1 complex, thus enhancing our understanding of the dynamic and intricate regulatory network during NLR-mediated immunity.

Unlike singleton NLRs, RRS1 and RPS4 form two-component immune complex that has been developed as a model for molecular understanding of paired NLR protein mechanisms. Despite its importance, the molecular mechanism involved in the control of complex formation and regulation remain poorly understood. In this complex, our previous studies revealed that RRS1 acts as a platform that enables the proper assembly of a functional RPS4/RRS1 complex and enhances the protein accumulation of RPS4^39,40^. Here, we revealed a reversible ubiquitination cycle in which the deubiquitinases UBP12 and UBP13 integrate with the E3 ligase RARE to coordinately modulate the complex turnover thorough (de)ubiquitinating the WRKY ID of RRS1. RARE directly interacts and ubiquitinates the integrated WRKY domain of RRS1, facilitating its proteasomal degradation. Since RRS1 stabilizes RPS4, RARE destabilization of RRS1 subsequently reduces RPS4 accumulation. Therefore, RARE impairs the complex abundance by promoting RRS1 and RPS4 degradation directly and indirectly, respectively. Since both RARE and AvrRps4 primarily associates with the WRKY ID, it is conceivable that AvrRps4 could compete with RARE for the same binding sites on RRS1, thus interfering with the interaction between RARE and RRS1 and antagonistically regulating RRS1 degradation during RRS1/RPS4-mediated defense. Based on the findings, we speculate that RARE keeps RRS1/RPS4 complex at a relatively low level in the resting state. However, upon perception of AvrRps4, AvrRps4 could down-regulate RARE-mediated degradation of RRS1. Consequently, the active state of RRS1/RPS4 is stabilized and accumulates to a sufficient level to induce a robust ETI response. Conversely, UBP12/UBP13 counteract RARE’s action, thereby preventing RRS1 over-ubiquitinated under resting state and safeguarding the complex. Our study reveals the delicate, reversible ubiquitination control of paired NLR complex levels that fine-tunes immunity, advancing understanding of paired NLR complex homeostatic regulation.

We further demonstrate reversible ubiquitination control of WRKY70/WRKY41 which are the putative virulence targets of AvrRps4 and PopP2 and homologous to the RRS1’s WRKY ID by RARE and UBP12/UBP13. Critically, we show this post-translational regulatory mechanism likely transfers from the WRKY proteins to the NLR-ID protein RRS1-R via integration of the WRKY domain, enabling homeostatic modulation of RRS1/RPS4-mediated immunity. Such evolutionary jumps likely synchronize the RRS1/RPS4 abundance with the WRKY70/WRKY41 levels, and possibly more other WRKY transcription factors. This aligns with the emerging view of unified plant immune system and concerted action of two sets of immune receptors since WRKY transcription factors are downstream components of PRRs required for basal immunity^46,47,49–52^.

Our finding of the transfer of reversible ubiquitination from WRKY70/WKYY41 to RRS1 through WRKY domain integration exposes an underexplored phenomenon with profound implications for protein regulation. It raises questions about the prevalence of this phenomenon and its impact on expanding organismal PTM regulatory network. Various extraneous domains like WRKY, zinc-finger BED and kinase domains have repeatedly integrated into NLR proteins across all plant lineages^8,9^. Non-integrated proteins harboring these homologous domains can undergo different types of PTMs such as phosphorylation, sumoylation or acetylation, suggesting the potential transfer of these modifications to regulate NLR-IDs’ function after domain integration, thus endowing NLR-ID fusion proteins with an additional layer of regulatory features. Similarly, the Src homology 2 (SH2) domain, a phosphotyrosine-binding domain indispensable for the receptor tyrosine kinase signalling pathways, represents another potential case of PTM transfer through domain shuffling. Rapid evolution of phosphotyrosine signaling by shuffling SH2 domains into various proteins with a range of biochemical properties including catalytic kinase domain endows SH2 domain with phospho-regulation by the Src family kinases^53,54^, echoing our findings.

In summary, our results are complementary to the previous work reporting RRS1/RPS4 complex is subjected to multilayered regulation, including phosphorylation, transcriptional regulation and alternative splicing^18,55,56^. This study has revealed a new layer of post-translational regulation by ubiquitination and deubiquitination of the ID of RRS1. The countering activities of RARE and UBP12/UBP13 antagonistically and dynamically control the turnover of RRS1/RPS4 complex to maintain immune homeostasis. Since the molecular mechanism of paired NLR protein complex regulation remains largely unknown, our study not only provides a novel mechanism for regulating paired NLR/NLR-ID complex homeostasis, but also presents an example in which transfer of PTM through domain integration, which allows the emergence of novel post-translational regulatory circuitries.

## Acknowledgments

We thank Dr. Xia. Cui from the Chinese Agricultural Academy of Sciences for providing *ubp12-2w* mutant seeds. We also thank Dr. Qian Chen (China Agricultural University) for her help with the in vitro ubiquitination assay. We thank Dr. Robert Heal and Dr. Joe Win from The Sainsbury Laboratory for the suggestions on manuscript improvement and phylogeny analysis, respectively. This research was supported by the National Natural Science Foundation of China (Grant No. 32100239, 32270307), Beijing NOVA Programme (Z211100002121066), the Marie Skłodowska-Curie scheme (H2020-MSCA-IF-2014-655295) and ERC Advanced Grant “Immunity by Pair Design.”

## Author Contributions

Conceptualization, H.G., J.H., and J.D.G.J.; Investigation, H.G., Z.C., J.H., J.L., and F.L.H.M.; Writing – Original Draft, H.G. and J.H.; Writing – Review & Editing, H.G., J.H., and J.D.G.J.; Funding Acquisition, H.G., and J.D.G.J.; Supervision, H.G., and J.D.G.J.

## Plant Materials and Growth Conditions

The wild-type *Arabidopsis thaliana* plants used in this study were the Columbia-0 (Col-0). The mutant *ubp12-2w* (GABI_742C10) seeds were provided by Dr. Xia. Cui (Chinese Academy of Agricultural Sciences, Beijing, China) and these mutants have been described by Cui et al (Cui et al., 2013). The *rare* mutant line (SALK_093498C) was ordered from the Arabidopsis Biological Resource Center (ABRC, Ohio State University, USA). *rrs1arrs1b* and *rrs1-3* (SALK_061602) has been described by Saucet SB, et al. (Saucet SB, 2015). Arabidopsis plants were grown in a growth room at 22 °C under a 14 h light/10 h dark photoperiod with light density of 100 μmol m^−2^ s^−1^. *Nicotiana benthamiana* (*Nb*) and *Nicotiana tabacum* (cultivar ‘Petite Gerard’) plants were grown at 24°C under long-day photoperiod (16 hr light/8 hr dark) with light density of 120 μmol m^−2^ s^−1^. Leaves of four- to five-week-old plants were used for *Agrobacterium*-mediated transient expression assays.

## Plasmid construction and Generation of Transgenic Plants

Plasmids were constructed using Golden Gate cloning as described previously (Ma et al., 2018; Guo et al., 2020). Briefly, to clone the full-length gene or individual fragment with different tags, all the full-length ‘‘domesticated’’ coding sequence (removal of internal BsaI/BbsI sites without change the encoded amino acid) gene/fragment without stop codon were cloned into the coding sequencing module level 0 vector (pICSL01005). The resulting constructs were assembled with different C-terminus tags into level 1 binary vector pICSL86977 with the CaMV35S promoter, Ocs terminator C-terminal epitope tags. For generation of N-terminal epitope tagging vector, the resulting constructs were assembled with different N-terminus tags into level 1 binary vector pICSL86900 with the CaMV35S promoter, Ocs terminator and N-terminal epitope tags. The C-terminus tags used in this study include C-TurboID(pICSL50040), C-3xFlag (pICSL50007), C-HA (pICSL50009), C-Myc (PICSL50010), C-GFP (PICSL50008), cYFP(pICSL50002) and nYFP(pICSL50003). The N-terminus tag used in this study includes N-Flag (pICSL30005). The primers used in this study are listed in Supplementary Table 2. For generation of transgenic plants, constructs were introduced into *Agrobacterium* strain GV3101 and subsequently transform Arabidopsis using the floral dip method. The harvested seeds from transformed Arabidopsis were resuspended in 0.1% agarose and detection of transgenic seeds using red fluorescent protein (DsRed) as a visible marker. Positive transformants were selected based on the seeds with bright red fluorescence under a hand-held lamp LUYOR-3415RG (Luyor Instrument, Shanghai).

## Protein extraction, immunoblot analysis and Co-IP assay

For Co-IP assays in *Nb* leaves was performed as described previously with minor modifications (Guo et al., 2020), *Nb* leaves were harvested 48 hours post agroinfiltratiion and ground in liquid nitrogen to a fine powder using a pestle and mortar. The powder was subsequently homogenized in an equal volume of ice-cold GTEN buffer (10% glycerol, 150 mM Tris-HCl pH 7.5, 1 mM EDTA, 150 mM NaCl) supplemented with 10 mM DTT, 0.2% (vol/vol) Nodinet-40, Anti-protease tablet (Complete EDTA-free Roche) and 2% PVPP (Polyvinylpolypyrrolidone) on a rotator for 20 min at 4°C. The lysates were then centrifuged at 5000 g for 30 min at 4°C, and the supernatant was filtered through 0.45 mm filters. The filtered protein extract was mixed with 3xSDS loading buffer (30% glycerol, 3.3% SDS, 94 mM Tris-HCl pH 6.8, 0.05% (vol/vol) bromophenol blue) supplemented with 10 mM DTT for input samples or for immunoblot analysis. Immunoprecipitations were perfomed using filtered extract incubated with Anti-DYKDDDDK G1 Affinity Resin (GenScript, L00432) or Anti-HA IP Resin (GenScript, L00777) at 4°C for 2 h with gentle shaking. After incubation, beads were collected via centrifugation at 7000 rpm for 30 seconds, washed three times in washing buffer (GTEN buffer supplemented with 10 mM DTT, 0.2% Nodinet-40 and Anti-protease tablet) and then suspended in 2XSDS loading buffer containing 10 mM DTT before boiling at 95°C for 5 min.

For Co-IP assays performed in Arabidopsis protoplasts, protoplasts co-transfected with the indicated plasmids were harvested after overnight incubation at 23°C and lysed with native extraction buffer (150 mM Tris-HCl, pH 7.5, 0.5% Nonidet P-40, 2 mM EDTA, 150 mM NaCl, 10% glycerol, 1 mM NaVO_3_, 5 mM NaF and 1Xprotease inhibitor cocktail (Roche)) by vigorous vortexing for 1 min. The lysates were centrifuged at 12,000g for 10 min at 4°C and supernatant was incubated with Anti-HA IP Resin (GenScript, L00777) for 2 h at 4°C with gentle mixing. The antibody-coupled agarose beads were collected and washed three times using native extraction buffer. The precipitated beads were then resuspened in 30 ml of 2X SDS loading buffer denatured for 10 min at 96 °C.

The co-immunooprecipated proteins were electrophoretically separated by SDS-PAGE and then transferred to Immobilon-P PVDF membranes (Merck Millipore) using a wet transfer apparatus (Bio-Rad). After blocking at room temperature for 1 h with 5% nonfat dry milk in 1× Tris-buffered saline containing 0.1% Tween 20, membrane was incubated with Horseradish Peroxidate (HRP) conjugated antibodies (Anti-FLAG M2, 1:10000 dilution, Sigma; Anti-GFP, 1:10000 dilution, Santa Cruz Biotechnology) in TBST supplemented with 5% milk by gentle agitation at room temperature for 1 h. The membrane was then washed 3 times with agitation for 5 minutes in TBST, and once in TBS (Tris-Buffered Saline). Proteins of interest were visualized by luminescence of HRP using the chemiluminescent substrate on a Tanon-5200 Chemiluminescent Imaging System (Tanon Science and Technology).

## Expression of recombinant protein, In Vitro Ubiquitination and Deubiquitination Assays

Constructs of GST-and HA-tagged RARE, GST tagged UBP12/UBP12 (m), GST tagged UBP13/UBP13 (m), His-and FLAG-tagged RRS1^WRKY^, His-and FLAG-tagged RRS1B^WRKY^, His-and FLAG-tagged WRKY33^WRKY^ were introduced into and expressed in Escherichia coli Transetta(DE3) for purification following the manufacturer’s instructions. The In vitro ubiquitination/deubiquitination assays were conducted following the procedure previously described (Jeong et al., 2017; Liu et al., 2022). Briefly, 2 μg ubiquitin, 50 ng AtUBA2, 100 ng AtUBC10, 500 ng RARE, and with or without 500 ng substrate proteins were added to 30 μl ubiquitination buffer (50 mM Tris-HCl, pH 7.5, 5 mM MgCl_2_, 2 mM ATP and 2 mM dithiothreitol). Reactions were incubated at 30°C for 2 h and the reaction mixtures were subsequently separated on SDS-PAGE followed by immunoblot analysis using anti-Ub, anti-FLAG, or anti-GST antibodies, respectively. For in vitro deubiquitination, after incubation with E1, E2 and E3 for 2 h at 30 L, Polyubiquitinated substrates proteins were purified and then added to deubiquitination buffer (50 mM Tris-HCl, pH 7.4, 150 mM NaCl, 5 mM MgCl_2_, and 2mM DTT) followed by the addition of GST-UBP12, GST-UBP12(m), GST-UBP13 or GST-UBP13(m). The deubiquitination reactions were terminated by adding 4XSDS loading buffer. The samples were taken at 0.5 and 1 h and subjected to immunoblot analysis using anti-Ub, anti-FLAG, or anti-GST antibodies, respectively.

## Bacterial growth assay

Pto DC3000(AvrRps4) or Pto DC3000 were grown on selective King’s B (KB) medium plates overnight. Bacterial cells were harvested, washed and resuspended in 10 mM MgCl_2_ at a concentration of OD_600_ = 0.0005 for infiltration. Leaves of five-week-old Arabidopsis were hand-infiltrated using a 1mL needleless syringe. The infected leaves are harvested at 0 and 3 days post inoculation to quantify bacterial colonies. 2 leaf discs from one plant (one leaf disc per leaf) were collected and were ground in 10 mM MgCl_2_, serially diluted, spotted on selective LB agar plates. The plates were incubated at 28°C for two days before counting. Statistical significance is indicated by letters (p < 0.01, one-way ANOVA followed by Tukey’s post hoc test).

## Subcellular Localization, BiFC assay in Arabidopsis protoplasts and confocal laser-scanning microscopy

BiFC analysis was performed in transient transfection assays as described previously (Guo et al., 2016). The combinations of plasmids containing N- and C-terminal YFP fusions were co-transformed into Arabidopsis protoplasts using PEG-calcium-mediated transformation. The transfected protoplasts were incubated overnight under dim light at 23°C before observation. Images were captured using a Leica TCS SP5 confocal fluorescence microscope. For GFP fluorescence, the excitation wavelength was 488 nm and the emission spectra were collected at 495 to 525 nm, respectively. For DAPI fluorescence, the excitation wavelength was 358 nm and the emission spectra were collected at 415 to 515 nm, respectively. For chlorophyll autofluorescence, the excitation wavelength was 488nm and emission spectra were collected at 650 to 750 nm, respectively.

## RNA extraction and Quantitative reverse transcription-PCR (qRT-PCR)

Total RNA was extracted from 3-week-old plants using TRIzol reagent and then treated with DNase I to remove DNA contamination. Reverse transcription was conducted using PrimeScript II 1st Strand cDNA Synthesis Kit (Takara) following manufacturer’s protocol. Quantitative RT-PCR were performed in the ABI QuantStudio 6 Flex Real-Time PCR System using the SYBR Premix ExTaq kit (Takara) in accordance with the manufacturer’s instructions. Three biological replicates were performed for each sample, and the expression levels were normalized to those of *UBQ10* using the comparative 2^-ΔΔCt^ method. Primers used for real-time PCR were listed in Supplementary Table 2.

## Proximity biotinylation, free biotin depletion and affinity purification by streptavidin-coated beads

One-week-old transgenic seedlings expressing TuboID-tagged bait proteins (0.4 g) were soaked in 50 μM biotin solution for 3h and quickly washed three times with ice cold water to stop labeling reaction and to deplete excess biotin. The seedlings were then dried with paper towels and subsequently ground into fine powder with liquid nitrogen. The sample powder was resuspended with protein extraction buffer (150 mM Tris-HCl, pH 7.5, 150mM NaCl, 0.5% Triton X-100, 0.5% Nonidet P-40, 0.5% sodium deoxycholate, Anti-protease tablet (Complete EDTA-free Roche)) with gentle mixing and then centrifuged at 12,000 g for 10 min. The resulting supernatant was then run through the Zeba™ Spin Desalting Column (Thermo Fisher Scientific, Catalog number 89893) according to manufacturer’s instruction to remove the excess biotin in lysates. The elution was then mixed with 50 μl of the equilibrated streptavidin agarose beads (Catalog number 20349, Thermo Fisher Scientific) and incubated overnight at 4 °C. The beads were subsequently washed 3 times with washing buffer (150 mM Tris-HCl, pH 7.5, 150 mM NaCl, 0.5% Nonidet P-40, Anti-protease tablet (Complete EDTA-free Roche)) and transferred to new low protein binding tube (Thermo Fisher Scientific). The biotinylated proteins were eluted by 30 μL 2×SDS sample buffer and detected by immunoblot analysis using streptavidin-HRP (Catalog Number. N100, Thermo Fisher Scientific)

## Phylogenetic analysis

To avoid long-branch attraction, the neighbour-joining method, rather than the maximum likelihood, was utilized to generate interspecies phylogenetic trees of RRS1, EARE and UBP12/13 under Poisson substitution model and uniform rates by MEGA 11^57^. On the other hand, the intraspecies phylogenetic tree of WRKY transcription factors was generated using the maximum-likelihood method under the JTT model and gamma distribution by IQ-TREE^58^.

For the phylogenetic tree of RRS1 orthologues, the protein sequences used include: *Arabidopsis thaliana*_RRS1-R (C4B7M5), *Arabidopsis lyrata*_RRS1 (XP_020871197), *Capsella rubella*_RRS1 (XP_006279894), *Camelina sativa*_RRS1 like (XP_010494612), *Raphanus sativus*_RRS1 (XP_018455889), *Brassica napus*_RRS1 like (XP_013734065), *Brassica rapa*_RRS1 (LOC103839134), *Brassica oleracea*_RRS1 (XP_013610929), *Arabidopsis thaliana*_CHS3 (NP_197291), *Arabidopsis thaliana*_CSA (NP_197290), *Arabidopsis thaliana*_RPS4 (NP_199338), *Theobroma cacao*_N (XP_017978840), *Eucalyptus grandis*_RPV (XP_010030110), *Vitis vinifera*_N (XP_010661240), *Solanum tuberosum*_NLR (KAH0750703), *Citrus sinensis*_Run1 like (XP_052293259), *Manihot esculenta*_NLR (KAG8633511), *Citrus clementina*_CICLE (ESR59447), *Carica papaya*_NLR (XP_021909738) and *Gossypium arboreum*_DCS1 (XP_017603119).

For the phylogenetic tree of RARE, the protein sequences used include: *Arabidopsis thaliana*_RARE (NP_001319035), *Capsella rubella*_RARE (XP_006307360), *Brassica rapa*_EARE (XP_009149355), *Citrus sinensis*_EARE (KAH9721972), *Manihot esculenta*_EARE (XP_021592540), *Vitis vinifera*_EARE (WKA08472), *Solanum tuberosum*_EARE (KAH0763893), *Oryza sativa*_EARE (KAF2939895), *Zea mays*_EARE (NP_001169658), *Amborella trichopoda*_EARE (XP_011623580), *Selaginella moellendorffii*_EARE (XP_024528416), *Marchantia polymorpha*_EARE (P888), *Chlamydomonas reinhardtii*_EARE (XP_042914639), *Dunaliella salina*_EARE (KAF5826418), *Arabidopsis thaliana*_PRT1 (NP_189124) and *Arabidopsis thaliana*_RMA3 (NP_194477).

For the phylogenetic tree of UBP12/13, the protein sequences used include: *Arabidopsis thaliana*_UBP12 (NP_568171), *Capsella rubella*_UBP12 (XP_006289281), *Brassica rapa*_UBP12 (XP_009122271), *Arabidopsis thaliana*_UBP13 (NP_001326292), *Capsella rubella*_UBP13 (XP_006299059), *Brassica rapa*_UBP13 (RID63933), *Citrus sinensis*_UBP12 (XP_006481665), *Manihot esculenta*_UBP12 (XP_021620269), *Vitis vinifera*_UBP12 (XP_002267555), *Solanum tuberosum*_UBP12 (XP_006339190), *Amborella trichopoda*_UBP12 (XP_020522273), *Oryza sativa*_UBP12 (XP_015617197), *Zea mays*_UBP12 (NP_001346364), *Selaginella moellendorffii*_UBP12 (XP_024540510), *Marchantia polymorpha*_UBP12 (P865), *Chlamydomonas reinhardtii*_UBP12 (XP_042922837), *Dunaliella salina*_UBP12 (KAF5839814), *Arabidopsis thaliana*_UBP26 (NP_001325635) and *Arabidopsis thaliana*_UBP16 (NP_567705).

To generate the phylogenetic tree of WRKY transcription factors, their WRKY domains were identified by the Conserved Domain Database^59^and the protein sequences of WRKY domains were subjected to phylogenetic analysis.

## LC-MS/MS Analysis

LC-MS/MS analysis was performed using an Orbitrap Fusion trihybrid mass spectrometer (Thermo Fisher Scientific) and a nanoflow-UHPLC system (Dionex Ultimate3000, Thermo Fisher Scientific) as previously described^60^ with the following modifications. Peptides were trapped to a reverse phase trap column (Acclaim PepMap C18, 5 μm, 100 μm×2 cm, Thermo Fisher Scientific) connected to an analytical column (Acclaim PepMan 100, C18 3 μm, 75 μm×50 cm, Thermo Fisher Scientific). Peptides were eluted in a sigmodial gradient of 3–65% acetonitrile in 0.1% formic acid (solvent B) over 90 min at a flow rate of 300 nl/min at 40 °C. The mass spectrometer was operated in positive ion mode with nano-electrospray ion source with an inner diameter of 0.02 mm fused silica emitter (New Objective). Voltage 2200 V was applied via platinum wire held in PEEK T-shaped coupling union with transfer capillary temperature set to 275 °C. The Orbitrap, MS scan resolution of 120,000 at 400 m/z, range 300 to 1800 m/z was used, and automatic gain control was set to 2 × 10^5^ and maximum inject time to 50 ms. In the linear ion trap, product ion spectra were triggered with a data-dependent acquisition method using “top speed” and “most intense ion” settings. The threshold for collision-induced dissociation (CID) and high energy collisional dissociation (HCD) was set using the Universal Method (above 100 counts, rapid scan rate, and maximum inject time to 10 ms). The selected precursor ions were fragmented sequentially in both the ion trap using CID and in the HCD cell. Dynamic exclusion was set to 30 s. Charge state allowed between +2 and +7 charge states to be selected for MS/MS fragmentation.

Peak lists in format of Mascot generic files (.mgf files) were prepared from raw data using MSConvert package (Matrix Science). Peak lists were searched on Mascot server version 2.4.1 (Matrix Science) against TAIR (version 11) database, a separate in-house constructs database and an in-house contaminants database. Tryptic peptides with up to two possible mis-cleavages and charge states +2, +3, +4 were allowed in the search. The following modifications were included in the search: oxidized Met, phosphorylation on Ser, Thr, Tyr as variable modifications, and carbamidomethylated Cys as a static modification. Data were searched with a monoisotopic precursor and fragment ions mass tolerance 10 ppm and 0.6 Da, respectively. Mascot results were combined in Scaffold version 4 (Proteome Software) and exported to Excel (Microsoft Office).

---

**Supplemental Figure 1.**
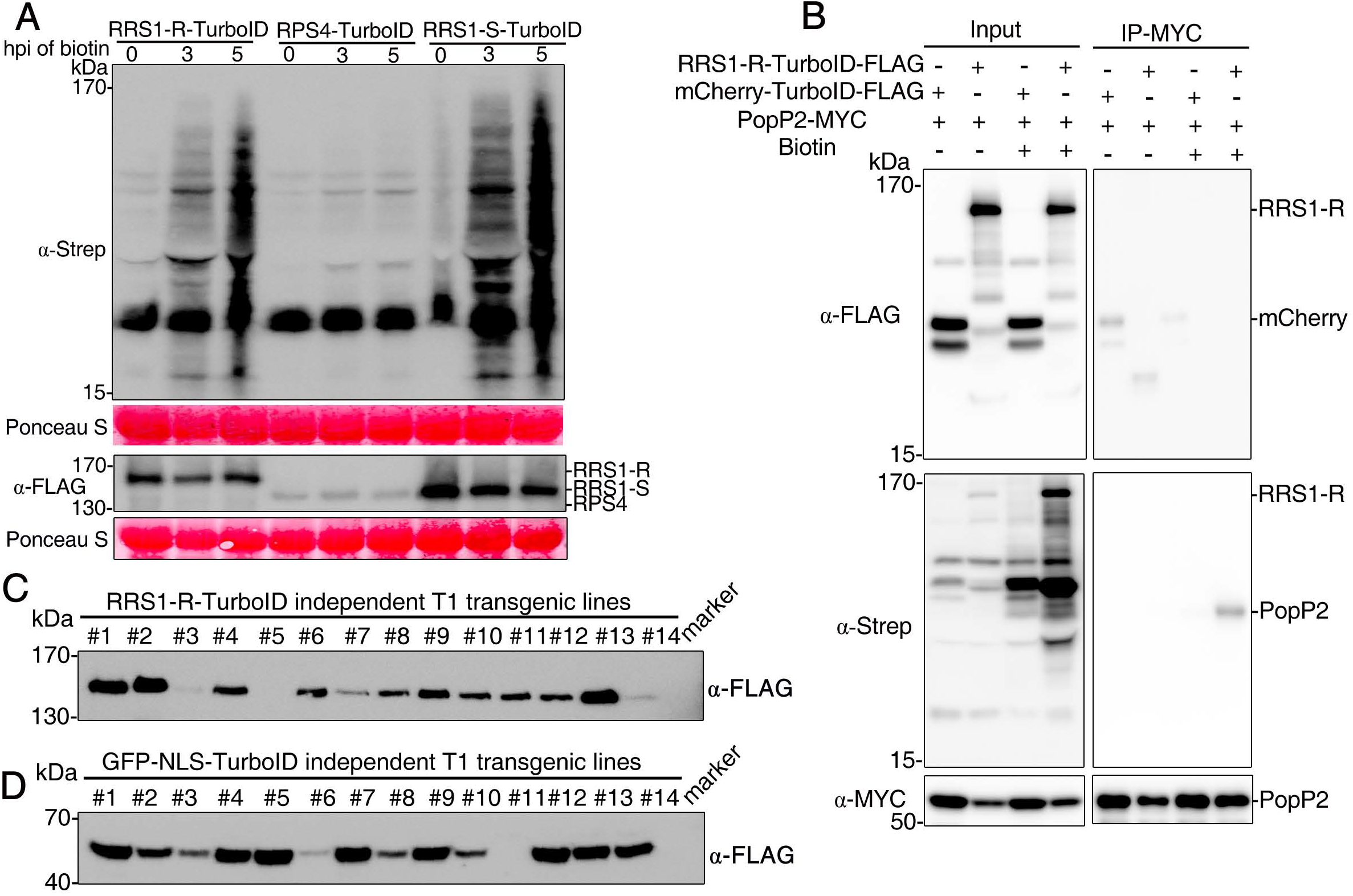
E**s**tablishment **of TurboID-mediated proximity labeling in Arabidopsis for identification of proximal proteins of RRS1/RPS4 immune receptor complex.** **(A)** RRS1-TurboID shows more efficient and clear biotinylation in *N. benthamiana*. *N. benthamiana* leaves were agroinfiltrated with the indicated construct and biotin was infiltrated into the previously agroinfiltrated leaves at 36 h post-agroinfiltration (hpi). Activity and expression of the TurboID fusion constructs were analyzed by immunoblot analysis with streptavidin-HRP and anti-FLAG antibodies, respectively. The experiment was repeated three times with similar results. Ponceau S staining for Rubisco are shown as loading controls. **(B)** TurboID-based analysis of PopP2 biotinylation by RRS1-R-TurboID in *N. benthamiana*. *N. benthamiana* leaves were agroinfiltrated with the indicated constructs combination and biotin was infiltrated into the previously agroinfiltrated leaves at 36 h post-agroinfiltration (hpi). Agrobacterial strains containing mCherry-TurboID infiltrated into the leaves served as a negative control. IP was carried out using leaf samples collected 3 h after biotin treatment with anti-MYC beads. The FLAG- and MYC-tagged proteins were detected using an anti-FLAG, anti-MYC and biotinylated proteins were detected using Streptavidin-HRP antibody, respectively. The experiment was repeated three times with similar results. **(C-D)**Plant lines generated for identification of proximal proteins of RRS1-R/RPS4 immune receptor complex. Protein accumulation of the RRS1-R-TurboID protein (C) and GFP-NLS-TurboID protein (D) in T1 lines used for the identification of proximal proteins of RRS1-R/RPS4 immune receptor complex. Line names are given on the top. Seeds harvested from *Agrobacterium*-transformed Arabidopsis were visual screened and seeds with bright red fluorescence are selected as the positive transformants. T1 transformants were transferred to fresh pots and leaves of 3-week-old Arabidopsis plants were harvested for protein extraction.

**Supplemental Figure 2.**
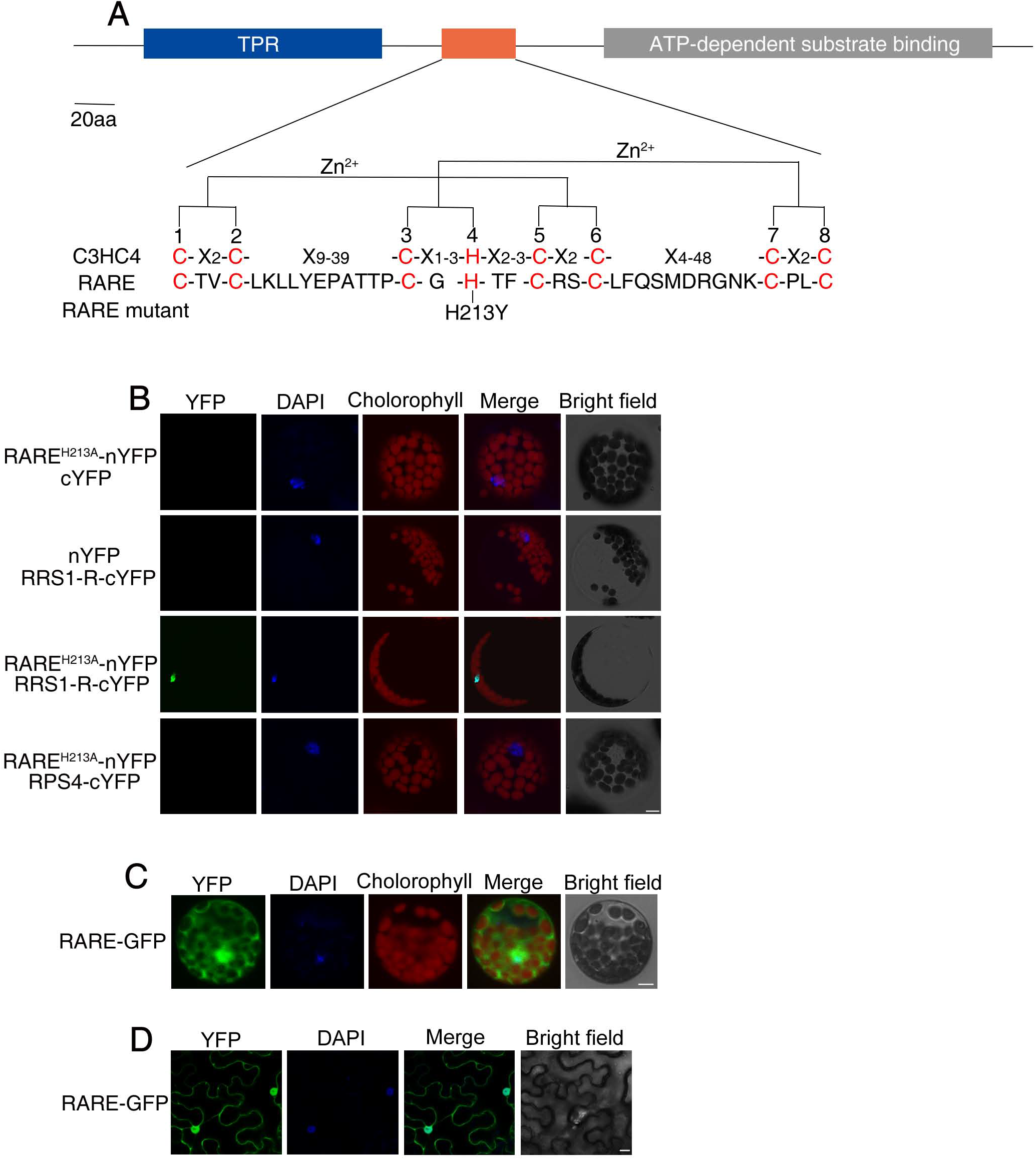
R**A**RE **is an E3 ubiquitin ligase that interacts directly with RRS1 and indirectly with RPS4 via RRS1.** **(A)** Domain architecture of RRS1-associated RING E3 ubiquitin ligase (RARE). The highly conserved RING finger domain is shown below the schematic diagram. The conserved cysteine (C) and histidine (H) residues involved in zinc binding are highlighted in red. Closed triangle indicates the mutation of conserved His residue. **(B)** BiFC analysis of the interaction of RARE with RRS1/RPS4 complex in Arabidopsis protoplasts isolated from *rrs1-3* plants. The indicated BiFC constructs were cotransfected into Arabidopsis protoplasts isolated from *rrs1-3* plants and the YFP fluorescence was visualized by confocal microscopy 16-20 h after transient expression. The positions of nuclei were shown by 4, 6-diamidino-2-phenylindole (DAPI) staining. Scale bar, 5 μm. **(C)** Subcellular localization of RARE-GFP in Arabidopsis mesophyll protoplasts. RARE-GFP (green) partially co-localized with DAPI signals (blue). Photographs were taken at 18 h after transformation using a confocal laser scanning microscope. Scale bar, 5 μm. **(D)** Subcellular localization of RARE-GFP in *Nb* leaves. RARE-GFP (green) partially co-localized with DAPI signals (blue). RARE-GFP fusion proteins were transiently expressed in *Nb* leaves and photographs were taken at 2 dpi. Scale bar, 10 μm.

**Supplemental Figure 3.**
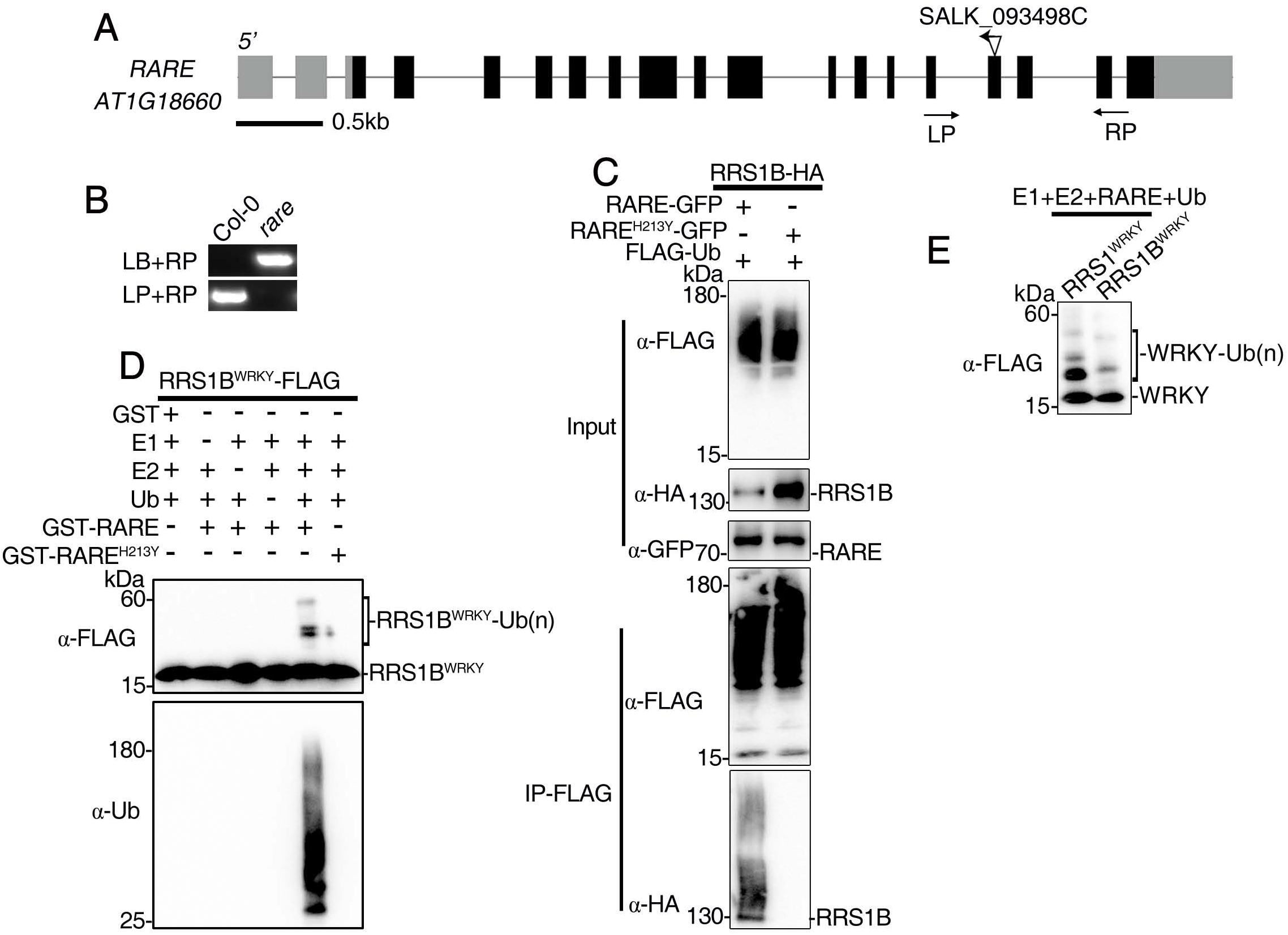
R**A**RE **is a functional E3 ligase and ubiquitinates the WRKY domain of RRS1.** (A) Schematic diagrams of the RARE gene structures with the T-DNA insertion sites. Black boxes indicate exons, gray boxes indicate untranslated regions, and lines indicate introns. The approximate positions of primers used for genotyping are indicated. LP and RP represent the forward primer and the reverse primer, respectively. (B) Identification of the homozygous T-DNA insertion mutant *rare*. T-DNA-specific primers and gene specific primers were used for genotyping. (C) Detection of ubiquitination of RRS1B by RARE in *Nb* leaves. HA-tagged RRS1B was co-expressed with GFP-tagged RARE or its mutant form RARE^H213Y^ in the presence or absence of FLAG-Ub in *Nb* leaves. The *Nb* leaves were pretreated with 50 μM MG132 for 6h before harvesting. Total ubiquitinated proteins were immunoprecipitated at 36h post-infiltration with anti-FLAG antibody, and ubiquitinated RRS1-R proteins were detected by immunoblotting with anti-HA antibody. (D) Ubiquitination of WRKY domain of RRS1B by RARE in vitro. FLAG-tagged WRKY domain of RRS1B was incubated with GST-RARE in ubiquitination assay buffer. Samples were resolved by SDS-PAGE and subjected to immunoblot analysis with anti-FLAG (top panel) or anti-Ub (bottom panel) antibody. Direct ubiquitination of WRKY domain was evident by higher molecular laddering detected by immunoblotting with anti-FLAG. (E) RARE less efficiently polyubiquitinated the WRKY domain of RRS1B compared to RRS1 in vitro. FLAG-tagged WRKY domain of RRS1 or RRS1B was incubated with GST-RARE in ubiquitination assay buffer. Direct ubiquitination of WRKY domain was evident by higher molecular laddering detected by immunoblotting with anti-FLAG antibody.

**Supplemental Figure 4.**
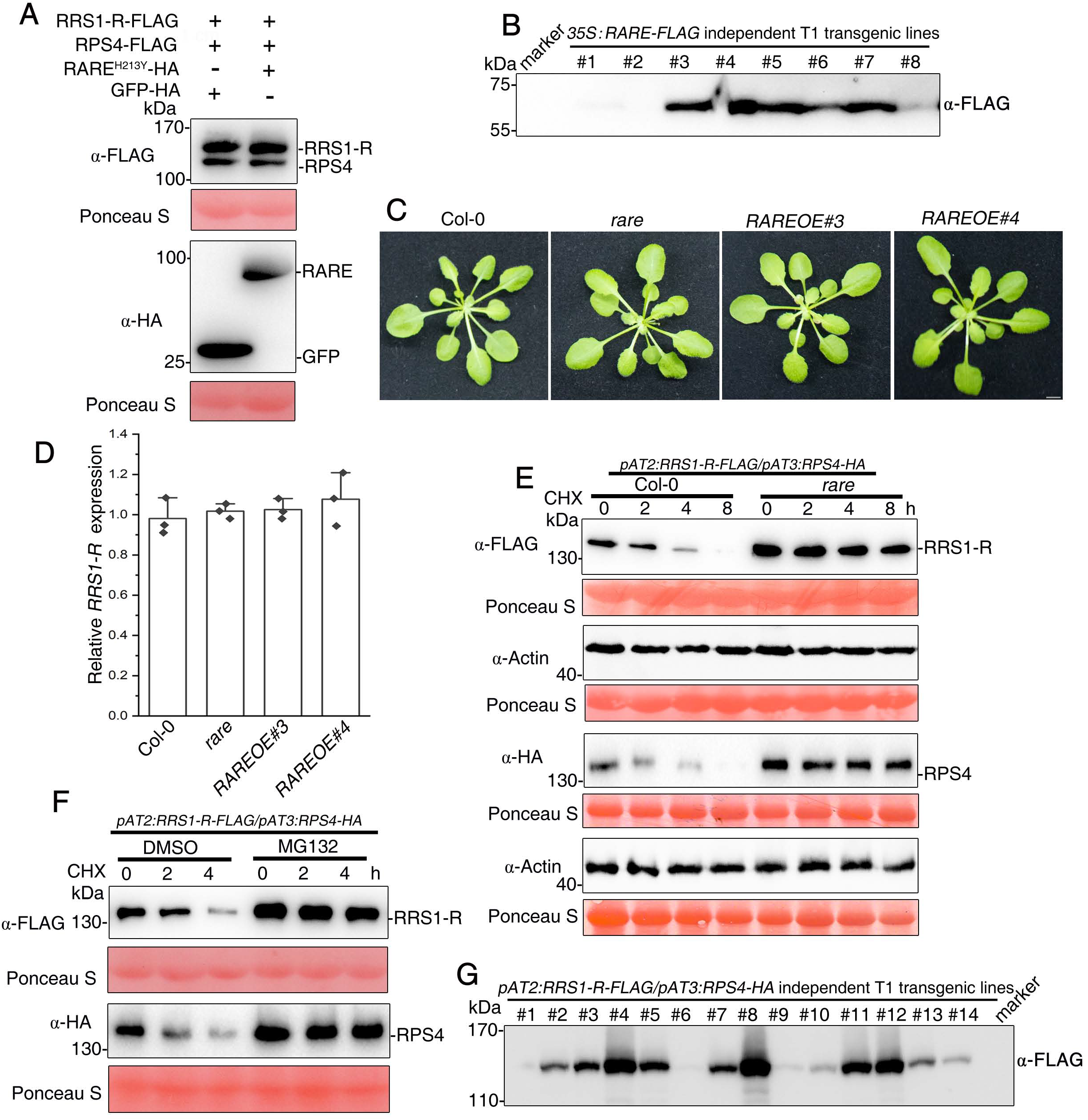
R**A**RE **destabilizes RRS1/RPS4 complex in a RRS1-dependent manner and negatively regulates RRS1/RPS4–mediated defense responses.** **(A)** RARE(H213Y) had little effect on the accumulation of RRS1/RPS4 complex. RRS1-R/RPS4 were transiently co-expressed with RARE(H213Y) in *Nb* at 2 dpi and then subjected to immunoblotting with anti-FLAG antibody for detecting RRS1-R/RPS4 or anti-HA antibody for detecting RARE. The HA-tagged GFP served as a negative control. **(B)** Protein accumulation of RARE in T1 transgenic lines. Protein accumulation of the RARE in FLAG-tagged ARAE transgenic plants. Leaves of 3-week-old T1 transformants were harvested for protein extraction followed by immunoblot analysis using anti-FLAG antibody. **(C)** The morphological phenotypes of WT (Col-0), *rare* and *RAREOE* plants. Four-week-old soil grown plants at 22℃ under long-day condition were photographed. Scale bar, 1 cm. **(D)** Real-time PCR analysis of *RRS1-R* transcript levels in *rare, RAREOE* and Col-0 plants. Total RNA was extracted from 7-day-old seedlings of Col-0, *rare* and *RARE* overexpression lines. The *RRS1-R* mRNA levels were measured by RT-qPCR. The relative transcript level of *RRS1-R* were normalized to those of *UBQ.* Value is mean ± SD of three independent replicates using independent seedling samples grown under the same conditions. Statistical significance compared with Col-0 was determined by Student’s t tests; ns, not significant. **(E)** Immunoblot analyses of RRS1-R and RPS4 protein turnover in Col-0 and *rare* mutant background. 10-days-old seedlings grown on MS medium were pretreated with 100 μM CHX for the indicated time before total protein was examined with immunoblot analysis. Total proteins were extracted from whole seedlings and subjected to immunoblot analysis using anti-FLAG antibody for detecting RRS1-R and anti-HA antibody for detecting RPS4, respectively. Actin was detected as a loading control. Note that the epitope-tagged RRS1-R/RPS4 transgenic line#2 was used to cross with *rare* and *RAREOE* transgenic line. **(F)** RRS1/RPS4 immune receptor complex undergoes proteasomal degradation. 10-days-old seedlings grown on MS medium were pretreated with 100 μM CHX for the indicated times in the presence or absence of 50 μM MG132. Total proteins were extracted from whole seedlings and subjected to immunoblot analysis with using anti-FLAG antibody for detecting RRS1-R and anti-HA antibody for detecting RPS4, respectively. Equal loading is shown by Ponceau S staining. **(G)** Protein accumulation of RRS1-R in T1 transgenic lines. Protein accumulation of the RRS1-R in FLAG-tagged RRS1-R transgenic plants. Leaves of 3-week-old T1 transformants were harvested for protein extraction followed by immunoblot analysis using anti-FLAG antibody.

**Supplemental Figure 5.**
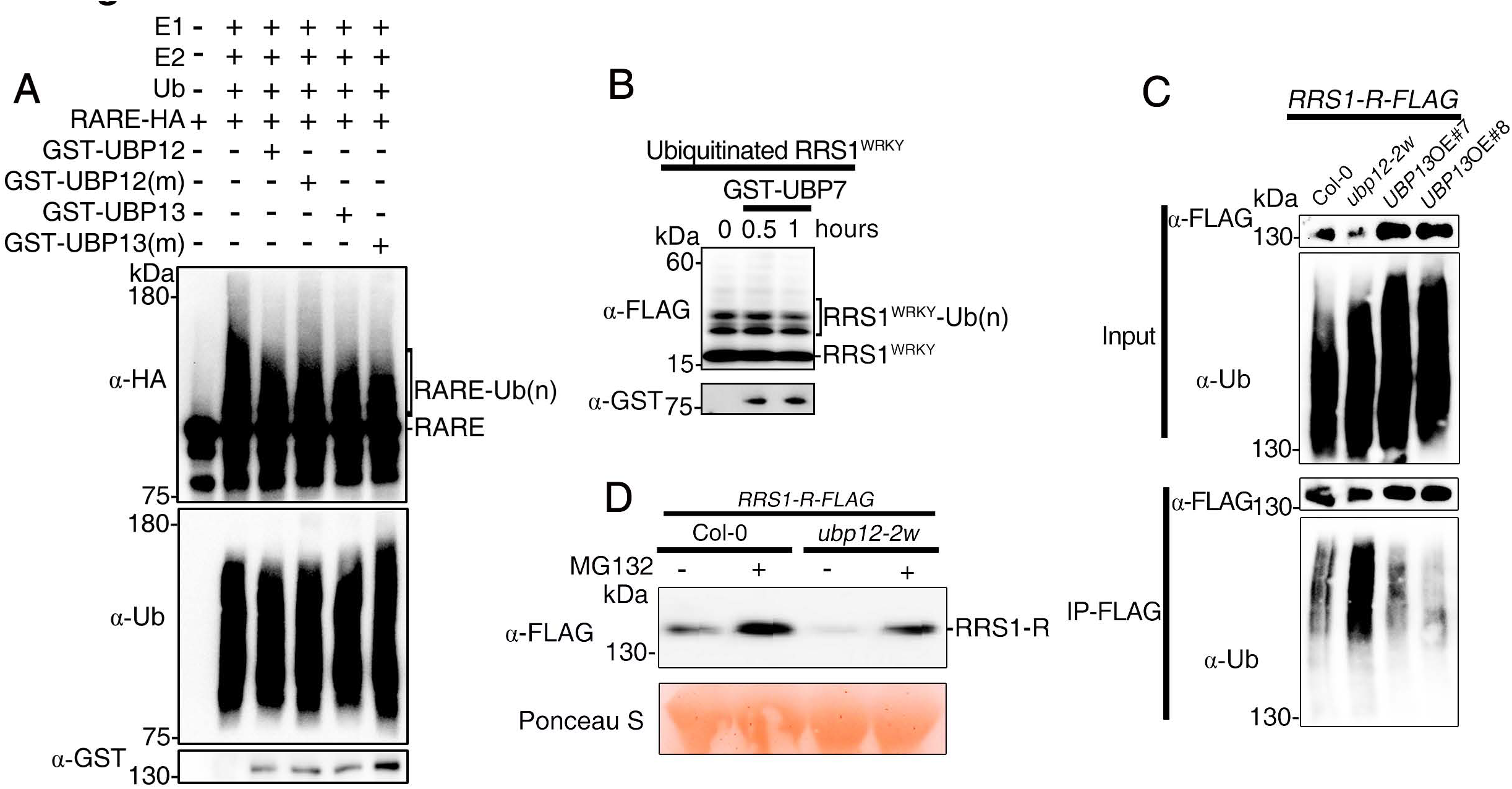
U**B**P12 **and UBP13 counteract the effect of RARE by removing ubiquitin from polyquibiquinated RRS1.** **(A)** UBP12 and UBP13 exhibit no activity toward polyubiquitinated RARE. Polyubiquitinated RARE produced by autoubiquitination was incubated with GST-UBP12 and GST-UBP13 for 2 h. Ubiquitinated RARE was detected by anti-HA (top panel) and anti-Ub antibodies (middle panel). UBP12 and UBP13 were detected by anti-GST antibody (bottom panel). **(B)** UBP7 has no activity toward polyubiquitinated WRKY domain of RRS1. Polyubiquitinated WRKY domain proteins was incubated with GST-UBP7 for 1 h. Ubiquitinated WRKY domain was detected by anti-FLAG (top panel). GST-UBP7 was detected with anti-GST antibody (bottom panel). **(C)** The polyubiquitination status of RRS1 in various genotypes. 10-days-old seedlings grown on MS medium were pretreated with 50 mM MG132 for 6 h. Protein extracts were immunoprecipitated with agarose-conjugated anti-FLAG antibody, followed by immunoblotting with anti-ubiquitin (top panel) and anti-FLAG antibodies (bottom panel). **(D)** Effect of MG132 on the RRS1 protein accumulation in *ubp12-2w* mutant background. 10-days-old seedlings grown on MS medium were supplied with or without 50 μM MG132 for 6h before harvesting. Total proteins were extracted from whole seedlings and subjected to immunoblot analysis using anti-FLAG antibody for detecting RRS1-R. Ponceau S staining of Rubisco indicates equal loading.

**Supplemental Figure 6.**
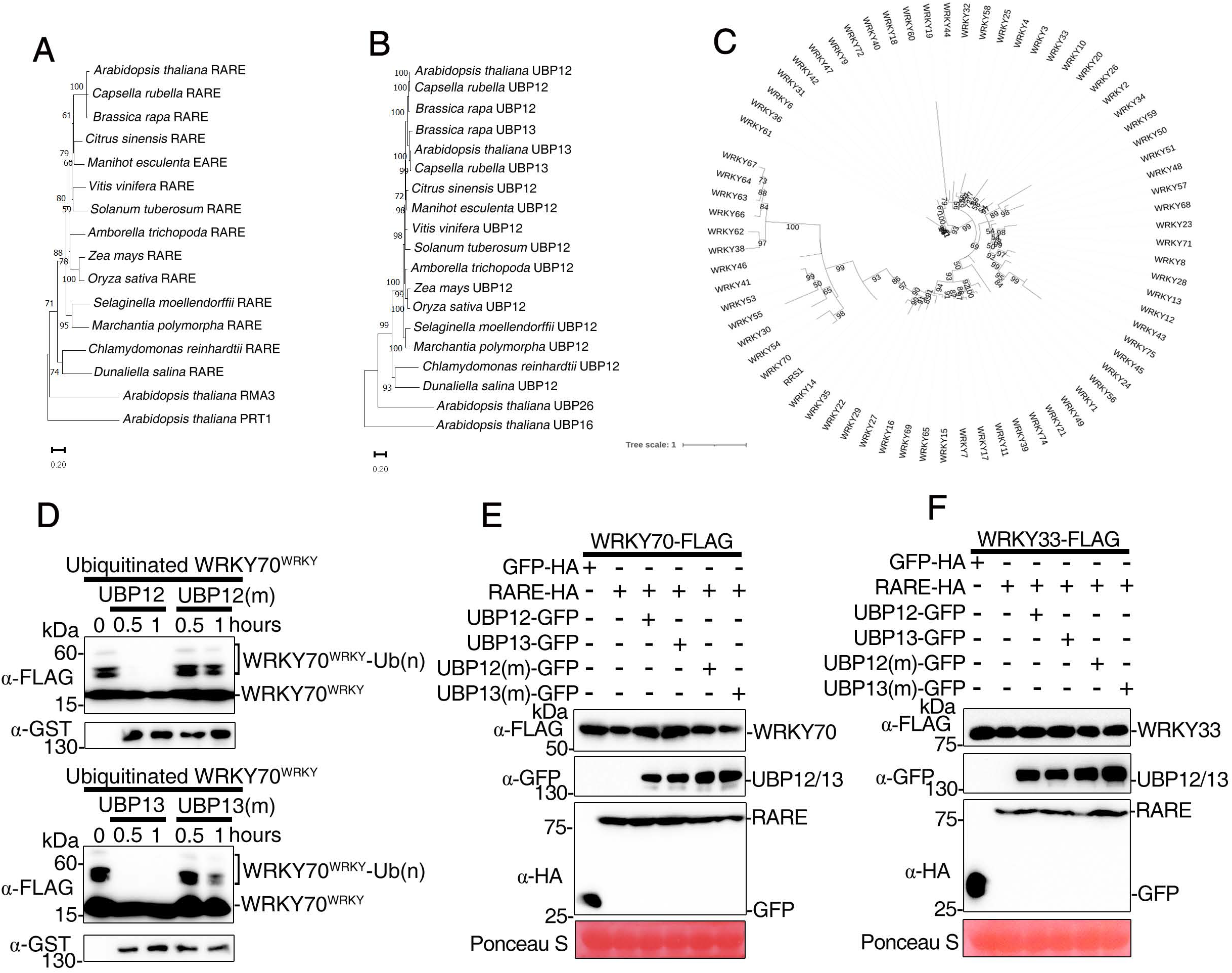
R**A**RE **and UBP12/UBP13 antagonistically regulate two WRKY transcription factors homologous to RRS1-R^WRKY^.** **(A and B)** The neighbour-joining phylogenetic trees of RARE orthologues (A) and UBP12/13 orthologues (B). The trees were generated based on full-length amino acid sequences. Only bootstrap values >50% within 1000 replicates are shown above branches. Ring-type E3 ligases ARI12 and BOI are outgroup for (A), whereas deubiquitinases UBP16 and UBP26 serve as outgroup for (B). **(C)** The maximum-likelihood phylogenetic tree of all 72 Arabidopsis WRKY transcription factors. The trees were generated based on amino acid sequences of their WRKY domains. Only bootstrap values >50% within 1000 replicates are shown above branches. **(D)** UBP12/UBP13 deubiquitinate polyubiquitinated WRKY domain of WRKY70 *in vitro*. Polyubiquitinated WRKY domain proteins produced by ubiquitination in the presence of E1(UBA2),E2 (UBC10), RARE and ubiquitin (Ub) were pulled down and then incubated with GST-UBP12, GST-UBP13, or their catalytically inactive variants for 0.5 or 1 hour. Ubiquitinated WRKY domain was detected by anti-FLAG (top panel). GST-UBP12, GST-UBP13 or the inactive variants were detected with anti-GST antibody (bottom panel). **(E)** UBP12/UBP13 antagonize RARE-mediated WRKY70 degradation. WRKY70 was co-expressed with RARE in *Nb* leaves in the presence or absence of UBP12/UBP13. GFP was coexpressed in the experiment as a control. **(F)** UBP12/UBP13 and RARE exhibited no effect on the accumulation of WRKY33. WRKY33 was co-expressed with RARE in *Nb* leaves in the presence or absence of UBP12/UBP13. GFP was coexpressed in the experiment as a control.

