## Supplementary Materials for "Reversible ubiquitination of integrated domain controls paired NLR immune receptor complex homeostasis"

**Supplementary Table 1.** A summary of proteins proximal to RRS1-R/RPS4 immune receptor complex identified using TurboID-based proximity labeling followed by mass spectrometry analysis(Please see attached Excel sheet).

**Supplementary Table 2.** Primers used for plasmid construction in this study.

| Purpose | Gene/Fragments | Vector | Name | Sequence (5’-3’) |
| --- | --- | --- | --- | --- |
| TurboID assay | RRS1 | pICSL01005 | RRS1-R-F | AATGAAGACATAATGACCAATTGTGAAAAGGAT |
|  |  |  | RRS1-R-R | AATGAAGACATCGAACCATAATCGAAGAATGTTGACC |
|  | GFP-NLS |  | GFP-S | AATGAAGACATAATGCCCAAGAAGAAGAGGAAGGTGATGGTGAGCAAGGGCGAG |
|  |  |  | GFP-R | AATGAAGACATCGAACCCTTGTACAGCTCGTCCATGCCG |
| Recombinant protein expression | UBP12 and UBP12(C208S) | pGEX-5X-1 | GST-UBP12-F | GGGATCCCCGAATTCATGACTATGATGACTCCGCC |
|  |  |  | GST-UBP12-R | ATGCGGCCGCTCGAGATTGTATATTTTTACCGGCT |
|  | UBP13 and UBP13(C207S) |  | GST-UBP13-F | GGGATCCCCGAATTCATGACTATGATGACTCCGCC |
|  |  |  | GST-UBP13-R | ATGCGGCCGCTCGAGATTGTATATTTTCACCGGCT |
|  | RARE and RARE(H213Y) |  | GST-RARE-F | GGGATCCCCGAATTCATGTCTAACGAAGACTCTCT |
|  |  |  | GST-RARE-R | ATGCGGCCGCTCGAGACAATCAAAAGAAAGGGACT |
|  | RRS1^WRKY^ | pET1a-HIS | His-RRS1^WRKY^-F | CTTTATTTTCAGGGGGACGTACCAAAAAAGGAGAA |
|  |  |  | His- RRS1^WRKY^-R | GCCACCGCCACCAGAAGCCTTGCGTTTAGTGGGCC |
|  | RRS1B^WRKY^ |  | His- RRS1B^WRKY^-F | CTTTATTTTCAGGGGTCAAAGAGTCGCCGAAAGAA |
|  |  |  | His- RRS1B^WRKY^-R | GCCACCGCCACCAGAATGGTTATGCTCAGA |
|  | WRKY41^WRKY^ |  | His- WRKY41^WRKY^ -F | AGAATCTTTATTTTCAGGGGCCAAAGTGGACAGAGCAAGT |
|  |  |  | His- WRKY41^WRKY^ -R | CCCGAGCCACCGCCACCAGATGCTACTGTGTGTTTTGGTT |
|  | WRKY70^WRKY^ |  | His- WRKY70^WRKY^ -F | AGAATCTTTATTTTCAGGGGCCCGTTAAGGGTAAAAGAGG |
|  |  |  | His- WRKY70^WRKY^ -R | CCCGAGCCACCGCCACCAGAACAAGTCTTGCTCTTGGGAG |
|  | WRKY33^WRKY^ |  | His- WRKY33^WRKY^ -F | AGAATCTTTATTTTCAGGGGGGGAATGGTGGTGGAAGCAA |
|  |  |  | His- WRKY33^WRKY^ -R | CCCGAGCCACCGCCACCAGAACCGCTACCACGAGCTGCAG |
|  | UBP7 | pGEX-5X-1 | GST-UBP7-F | GGGATCCCCGAATTCATGGTGATCAAACCCGACCC |
|  |  |  | GST-UBP7-R | ATGCGGCCGCTCGAGCATGGAGATGAGACGGGCCT |
| Construction of transgenic plants | *RAREOE* | pCBFR C-FLAG | RAREOE-FLAG-F | CTATTCTAGTCGACCTGCAGGGCCATTACGGCCATGTCTAACGAAGACTCTCT |
|  |  |  | RAREOE-FLAG-R | CCCGAGCCACCGCCACCAGAGGCCGAGGCGGCCCCACAATCAAAAGAAAGGGACT |
|  | *UBP12OE* | pCAMBIA1300 | UBP12OE-GFP-F | ACGGGGGACGAGCTCGGTACCATGACTATGATGACTCCGCC |
|  |  |  | UBP12OE-GFP-R | TCTCCTTTGCCCATGTCGACATTGTATATTTTTACCGGCT |
|  | *UBP13OE* |  | UBP13OE-GFP-F | ACGGGGGACGAGCTCGGTACCATGACTATGATGACTCCGCC |
|  |  |  | UBP13OE-GFP-R | TTTGCCCATGTCGACATTGTATATTTTCACCGGCT |
| Co-immunoprecipitation Assay | Ubiquitin14 | pICSL01005 | Ubiquitin14-F | AATGAAGACATAATGCAGATCTTTGTTAAGACTC |
|  |  |  | Ubiquitin14-R | AATGAAGACATCGAACCACCACCACGGAGCCTGAGAAC |
| Mutants confirmation |  |  | GABI_742C10LP | CCTGGAAGCGATTATAAACAACAT |
|  | *ubp12-2w* |  |  |  |
|  |  |  | GABI_742C10RP | TAGGCTGCACCTTGTATTTTTCTT |
|  | *rare* |  | SALK_119141CLP | TCCCCTGTCAGAAGTTGTCTC |
|  |  |  | SALK_119141CRP | TCGTACATCTGCTTTCCCTTG |
| qPCR | *UBQ10* |  | UBQ10-F | CACACTCCACTTGGTCTTGCGT |
|  |  |  | UBQ10-R | TGGTCTTTCCGGTGAGAGTCTTCA |
|  | *PR1* |  | PR1-F | AGGCAACTGCAGACTCATAC |
|  |  |  | PR1-R | TTGTTACACCTCACTTTGGC |
|  | *PR2* |  | PR2-F | GCTTCCTTCTTCAACCACACAGC |
|  |  |  | PR2-R | CGTTGATGTACCGGAATCTGAC |
| Primers used to generate mutations | UBP12C208S |  | UBP12C208S-F | AGGTGCAACATCCTACATGAATTC |
|  |  |  | UBP12C208S-R | GAATTCATGTAGGATGTTGCACCT |
|  | UBP13C207S |  | UBP13C207S-F | GGTGCTACCTCTTACATGAATTCTC |
|  |  |  | UBP13C207S-R | GAGAATTCATGTAAGAGGTAGCACC |
|  | RARE(H213Y) |  | RARE(H213Y)-F | CTCCATGTGGGTATACATTCTGCCG |
|  |  |  | RARE(H213Y)-R | CGGCAGAATGTATACCCACATGGAG |
